# POINT Technology Illuminates the Processing of Polymerase-Associated Intact Nascent Transcripts

**DOI:** 10.1101/2020.11.09.374108

**Authors:** Rui Sousa-Luis, Gwendal Dujardin, Inna Zukher, Hiroshi Kimura, Maria Carmo-Fonseca, Nick J Proudfoot, Takayuki Nojima

## Abstract

Mammalian chromatin is the site of both RNA polymerase II (Pol II) transcription and coupled RNA processing. However, molecular details of such co-transcriptional mechanisms remain obscure, partly due to technical limitations in purifying authentic nascent transcripts. We present a new approach to purify and profile nascent RNA, called Polymerase Intact Nascent Transcript (POINT) technology. This three-pronged methodology maps nascent RNA 5’ends (POINT-5), establishes the kinetics of co-transcriptional splicing patterns (POINT-nano) and profiles whole transcription units (POINT-seq). In particular we show by depletion of the nuclear exonuclease Xrn2 that this activity acts selectively on cleaved 5’P-RNA at polyadenylation sites. Furthermore POINT-nano reveals that splicing occurs either immediately after splice site transcription or is delayed until Pol II transcribes downstream sequences. Finally, we connect RNA cleavage and splicing with either premature or full-length transcript termination. We anticipate that POINT technology will afford full dissection of the complexity of co-transcriptional RNA processing.

**HIGHLIGHTS:**

1. POINT methodology dissects intact nascent RNA processing
2. Specificity of Xrn2 exonuclease in co-transcriptional RNA degradation
3. Splicing suppresses Xrn2-dependent premature termination
4. Different kinetic classes of co-transcriptional splicing in human genes

## INTRODUCTION

Transcripts synthesized by eukaryotic RNA polymerase II (Pol II) are processed during transcription to gain functionality and control RNA stability. Multiple enzymatic reactions are involved in chromatin associated RNA processing, such as pre-mRNA splicing, transcript cleavage and selective degradation (Cramer, 2019; Desterro et al., 2020; Peck et al., 2019). First a Cap structure (7meGppp) is added to the transcript 5’end soon after its exit from the Pol II complex. This initial step defines all Pol II transcription and is ultimately required for efficient mRNA export and protein translation. As Pol II transcribes into the gene body, intronic RNA is rapidly removed by splicing. This involves assembly of the megadalton spliceosome complex, beginning with recognition of the intron 5’ splice site (SS) by U1 snRNA-protein complex (U1 snRNP). Once Pol II reaches the intron 3’end, U2 snRNP identifies the intronic branch point and 3’SS followed by assemblage of a further complex set of U snRNPs (U4, U5 and U6) and other associated protein factors. The spliceosome so formed, reorganizes the intron into a ribozyme-like structure, leading to intron excision and ligation of upstream to downstream exons. The above outline of the splicing mechanism implies an intron definition mechanism, where the appearance of intronic splice signals leads to a stepwise assembly of the spliceosome. However, in higher eukaryotes where exons are generally much shorter than their adjacent introns, it is thought that splicing factors such as SR proteins initially bind to and define functional exons, known as the exon definition mechanism (Ule and Blencowe, 2019). These will in turn facilitate recruitment of U2 snRNP to the 3’SS and U1 snRNP to the 5’SS leading to spliceosome formation. At the gene end (transcript end site or TES), specific polyadenylation signals (PAS) are recognized by the cleavage and polyadenylation (CPA) complex. An endonuclease CPSF73 within CPA cleaves the nascent RNA at the PAS and this is coupled with upstream RNA polyadenylation by polyA polymerase (Mandel et al., 2006). This critical RNA processing step promotes release of the mature mRNA from chromatin into the nucleoplasm and its subsequent transport to cytoplasmic ribosomes. The downstream RNA cleavage product is then degraded by the nuclear 5’-3’exonuclease, Xrn2 which follows behind the elongating Pol II and induces transcription termination upon reaching the Pol II complex (Proudfoot, 2016). Nascent transcript cleavage also occurs in other transcript regions such as at pre-microRNA (miRNA) sequences. Here hairpin structures are recognized by the double strand (ds)RNA specific endonuclease complex called the Microprocessor, comprising a dsRNA binding protein DGCR8 and dsRNA specific endonuclease DROSHA (Ha and Kim, 2014). Notably dsRNA cleavage in exons, but not in introns, promotes Pol II transcription termination. Likely splicing impedes termination otherwise induced by DROSHA cleavage (Dhir et al., 2015).

These RNA processing and cleavage events are tightly regulated during synthesis of all Pol II transcription units (TU) which begins at the TSS (transcription start site) and ends at the TES. Several nascent RNA analyses have contributed to our understanding of how these RNA processing events are coupled to transcription (Stark et al., 2019). For example, the method of transient transcript-sequencing (TT-seq) provides information on the extent Pol II TUs (Schwalb et al., 2016). We have also employed mammalian native elongating transcript-sequencing (mNET-seq) and with this methodology demonstrated that the S5P phospho isoform of Pol II C-terminal domain (CTD) is associated with Pol II pausing on spliced exons and in the recruitment of catalytic spliceosome (Nojima et al., 2015; Nojima et al., 2018a; Schlackow et al., 2017). Furthermore, mNET-seq revealed the kinetics of pre-miRNA cleavage and characterized Pol II pausing at the TES. A critical limitation of both these methodologies is the restricted length of nascent transcript reads that can be obtained. This is limited to less than 150 nt in most cases due to RNA fragmentation in the protocols employed, coupled with size limitations set by the Illumina sequencing platform. Recently long read sequencing using Oxford Nanopore Technology (ONT) has been employed to sequence 4sU metabolically labelled RNA as a way to dissect co-transcriptional splicing (Drexler et al., 2020). However, this approach is marred by the presence of mature 4sU labelled mRNA contamination that may confound the view that these data reflect largely nascent transcription.

We here describe new technology to dissect the complex Pol II transcription cycle by analysing intact nascent RNA directly purified from elongating Pol II. We name this technology <u>Po</u>lymerase <u>I</u>ntact <u>N</u>ascent <u>T</u>ranscript (POINT). Our POINT methodology employs both Illumina and ONT sequencing platforms. For Illumina we use *in vitro* fragmented RNA purified from immunoprecipitated (IP) Pol II elongation complexes to profile the nascent RNA across the whole TU (POINT-seq). We also employ unfragmented RNA in a 5’RACE-template switching protocol that maps nascent RNA 5’ends at single nucleotide resolution (POINT-5). Notably our POINT-5 method precisely maps and distinguishes TSS and RNA cleavage sites on pre-mRNA, pre-miRNA, histone and U snRNA genes. Xrn2-dependent RNA degradation at pre-mRNA TES is also detected by this technology. We further employ the ONT direct cDNA sequencing platform to characterise nascent RNA isolated by POINT technology (POINT-nano). This allows us to directly analyse the kinetics of splicing and CPA-mediated RNA cleavage. In effect our data not only provides new information on co-transcriptional RNA processing, but in so doing reveals the power of POINT technology as a way to unravel the complexity of transcriptional regulation and its cross-talk with RNA processing in eukaryotic chromatin.

## RESULTS

### Development of POINT technology

Analysing authentic nascent RNA is critical to fully comprehend the mechanisms of Pol II transcription and associated transcript processing. We have previously employed mNET-seq to map the 3’ ends of nascent RNA, which involves chromatin isolation and its digestion by micrococcal nuclease (MNase) to nucleosomal sized fragments. These are IPed with Pol II antibodies to yield RNA from within the Pol II active site. This short RNA fraction (20-100 nt) is detected by RNA linker ligation onto the transcript 3’end, followed by cDNA sequencing. mNET-seq can also identify the 3’ends of RNA processing intermediates within co-IPed spliceosome and microprocessor complexes (Nojima et al., 2015). Since mNET-seq generates only short RNA sequence information, this limits analysis of co-transcriptional RNA processing, due to lack of information on the exact position where transcription starts, as well as on the continuity of the RNA molecule. To overcome these size limitations, we have developed a different procedure to isolate intact nascent transcript from the 5’cap site through to its 3’end within the Pol II active site; POINT technology (Figure 1A). In outline, nuclear derived chromatin is purified in the presence of both 1M urea and 3% Empigen detergent. This treatment more fully denatures chromatin and so allows for efficient DNA specific digestion by DNase which unlike MNase cannot access DNA in standard chromatin preparations. Pol II with its associated nascent transcript is thereby solubilised (Figure S1A). Notably while the chromatin associated DNA is fully digested to nucleosome size fragments within 4 min, a longer digestion period (12 min) more completely digests all DNA outside of the Pol II elongation complex. In this way potential background proteins and RNA non-specifically associated with DNA, external to Pol II is eliminated (Figure S1B and S1C). Following DNase digestion, intact nascent RNA ending within the Pol II active site is isolated by immunoprecipitation (IP) in NET-2 buffer with additional 3% Empigen to remove all steady state mRNA and rRNA from the Pol II IP fraction (Figure S1D). The IPed RNA which has a size range from a short length to greater than 6,000 nt is either fragmented and sequenced (POINT-seq) or directly subjected to 5’RACE and template switching (POINT-5) using the Illumina strand-specific RNA-seq platform (Figure 1A).

**Figure 1.**
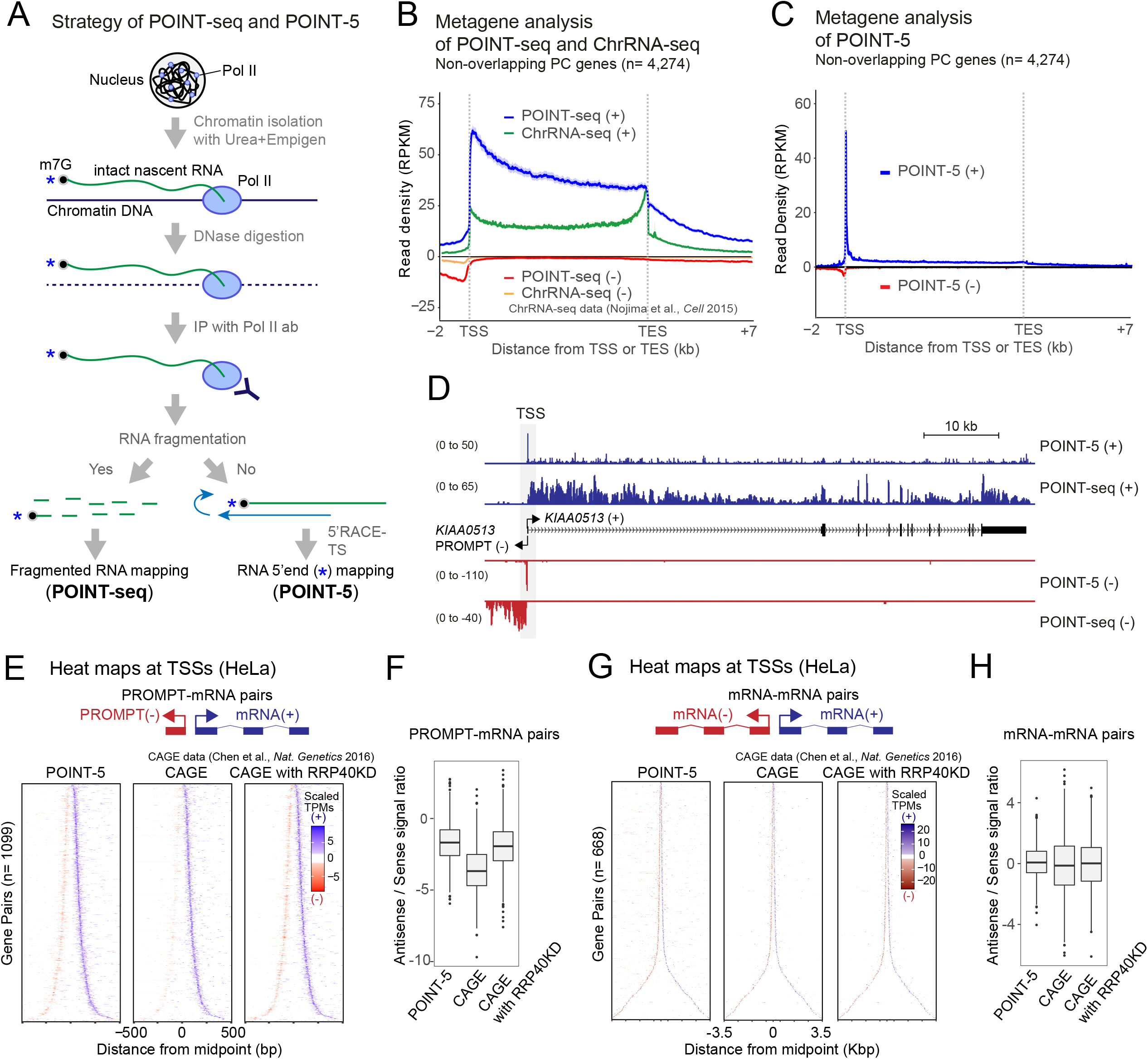
Development of POINT-seq and POINT-5 methodology. (A) Strategy of POINT technology. Chromatin fraction isolated with 1M Urea and 3% Empigen from nuclear preparations and digested with DNase. Intact nascent RNA IPed with Pol II antibody and purified. For POINT-seq, RNA fragmented during library prep. For POINT-5, non-fragmentated RNA reverse-transcribed with random hexamer and template switching during library prep. Both POINT-seq and POINT-5 libraries were Illumina sequenced. (B) Metagene of POINT-seq and ChrRNA-seq signals in normalized transcription unit from TSS −2kbp to TES +7kbp. POINT-seq profiles at (+) and (−) stands are shown in blue and red, respectively. Published Chromatin RNA (ChrRNA)-seq profiles at (+) and (−) stands are shown in green and orange. (C) Metagene of POINT-5 signals in normalized transcription unit from TSS −2kbp to TES +7kbp. POINT-5 profiles at (+) and (−) stands are shown in blue and red. (D) *KIAA0513* as example of POINT-5 and POINT-seq profiles. (E) Heat maps of POINT-5 and CAGE (untreated or RRP40KD) signals for PROMPT-mRNA pairs (n=1,099) at TSS. (F) Ratio of sense (+) and antisense (−) POINT-5 and CAGE (untreated or RRP40KD) signals at TSS. (G) Heat maps of POINT-5 and CAGE (untreated or RRP40KD) signals for mRNA-mRNA pairs (n=668) at TSS. (H) Ratio of sense (+) and antisense (−) POINT-5 and CAGE (untreated or RRP40KD) signals at TSS.

To validate our POINT methodology as a strategy to purify exclusively nascent RNA, we compared POINT-seq data with chromatin-bond RNA (ChrRNA-seq) data from HeLa cells (Nojima et al., 2015). ChrRNA-seq meta-profiles show highest signal near the TES, followed by a sharp drop indicative of substantial contamination by mRNA that have been cleaved at the TES and released into the nucleoplasm. In contrast the POINT-seq profiles detected highest signal over the TSS that gradually declined across the TU as well as higher signals in antisense and termination regions (Figure 1B). These different profiles clearly underline the predominantly nascent transcription detected by POINT methodology. Additionally, transcriptional inhibition by 4 hr treatment with DRB dramatically reduced POINT-seq signals over protein coding genes including *TARS* (Figure S1E). Importantly mature, exonic RNA was undetectable in POINT-seq from DRB-treated cells. These results confirm that POINT-seq profiles reflect highly purified nascent RNA. We also note that our POINT-seq data is reproducible between three biological replicates (Figure S1F).

POINT-5 profiles allowed detection of all 5’ends of non-overlapping protein coding genes expressed in HeLa cells, indicative of active TSS (Figure 1C) and show overall high data reproducibility (Figure S1G). We also employed POINT-5 to characterise TSS from divergent TUs and show that this method effectively maps the 5’ends of promoter upstream transcripts (PROMPT) as shown for *KIAA0513* (Figure 1D) as well as for various divergent mRNA-mRNA and eRNA-eRNA pairs (Figure S1H-S1J). Next, we compared POINT-5 to published Cap-Analysis of Gene Expression (CAGE) (Chen et al., 2016). CAGE detected robust signals for mRNA but little PROMPT signal (Figure 1E). This increased upon depletion of the exosome component RRP40 (Figures 1E and 1F) as expected since PROMPTs are known to be degraded by the exosome (Andersson and Sandelin, 2020; Preker et al., 2008; Schlackow et al., 2017). In contrast RRP40 depletion had no effect on CAGE signal from divergent mRNA (Figures 1G and 1H). Notably POINT-5 detected significant levels of PROMPT signal without exosome depletion (Figures 1E and 1F). These comparative analyses of POINT-5 versus CAGE emphasise the remarkably nascent nature of the POINT-5 data and confirm that this methodology provides a reliable approach to detect all newly synthesized, capped Pol II transcripts.

### POINT-5 defines co-transcriptional RNA cleavage sites

Nascent RNA can be cleaved during transcription by complexes containing endonucleases such as the Microprocessor that specifically cleaves dsRNA at the base of RNA stem-loop structures as formed for pre-miRNA (Morlando et al., 2008; Nojima et al., 2015; Pawlicki and Steitz, 2008). Also, the U7 snRNA-CPA complex cleaves histone transcripts following their downstream stem-loop (SL) structure (Marzluff and Koreski, 2017), while the Integrator complex is known to terminate U snRNA gene transcription (Chen and Wagner, 2010). Each of complex possess endonuclease activity, thought to act co-transcriptionally. Indeed, we show here that POINT-5 methodology can map co-transcriptional RNA cleavage sites at single nucleotide resolution, as with TSS, by detecting newly synthesised 5’ends of nascent RNA associated with the Pol II active site (Figure 2A).

**Figure 2.**
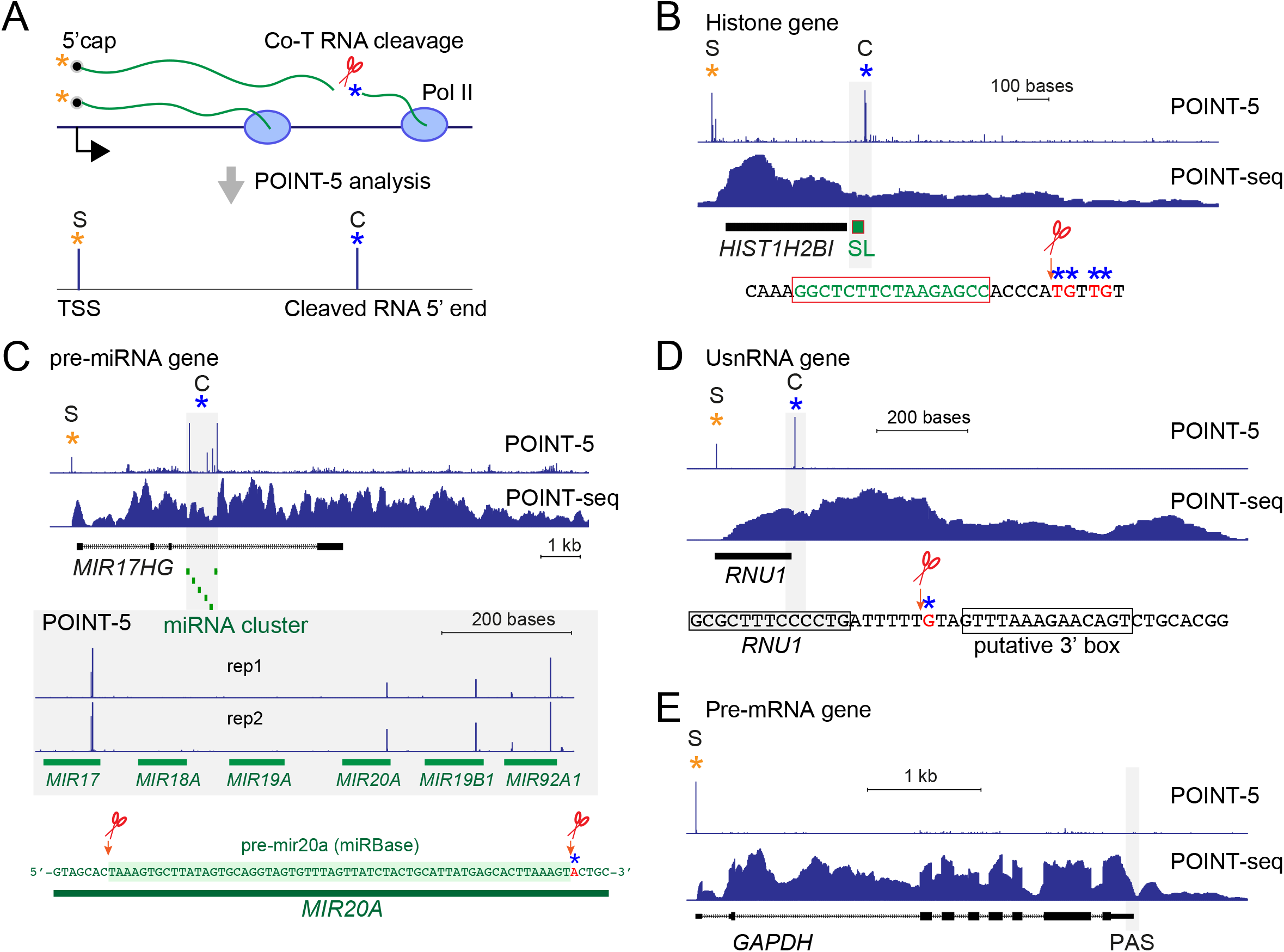
Detection nascent RNA 5’ ends by POINT-5. (A) Schematic diagram of POINT-5 analysis detecting TSS (S, orange asterisk) and co-transcriptional (Co-T) RNA cleavage site (C, blue asterisk). (B) Histone gene *HIST1H2BI* as example of POINT-5 and POINT-seq profiles. Stem loop (SL) structure highlighted in green. (C) Pre-miRNA gene, *MIR17HG* as example of POINT-5 and POINT-seq profiles. Cluster of miRNA are highlighted in green. POINT-5 peak was detected near 3’end of *MIR20A* as previously reported in miRBase. (D) U snRNA gene, *RNU1* as example of POINT-5 and POINT-seq profiles. Co-T RNA cleavage was detected upstream of putative 3’box. (E) Pre-mRNA gene, *GAPDH* as example of POINT-5 and POINT-seq profiles. No POINT-5 peak detected at TES.

We first applied POINT-5 technology to define endonuclease cleavage positions across a human histone gene locus containing multiple histone genes (Figure S2A). Clear peaks were detected at either end of each histone gene, one at the T<u>S</u>S (S peak) and another resulting from endonuclease <u>c</u>leavage (C peak). In particular for *HIST1H2BI*, the C peak showed local heterogeneity over several nucleotides, downstream from the SL structure that defines the 3’end of histone mRNA (Figure 2B). This is bound by SLBP, required both for histone transcript 3’end processing and subsequent histone mRNA translation. POINT-5 analysis also detected significant peaks indicative of DROSHA cleavage activity over intronic MIR26B in *CTDSP1* and lncRNA derived MIR193A and MIR365B (Figures S2B and S2C). Notably 3’ends of the co-transcriptionally cleaved pre-miRNA cluster MIR17-92A embedded within the lncRNA *MIR17HG* were previously detected by mNET-seq (Nojima et al., 2015). Our POINT-5 method instead maps the 5’end of these DROSHA cleaved RNA associated with Pol II. Published ENCODE databases profile expression of all mature small RNA derived from this cluster, while POINT-5 and mNET-seq data only detects co-transcriptionally cleaved 5’ and 3’ends for MIR17, 20a, 19b-1 and 92a-1, but not MIR18a and 19a (Figure 2C). These results underline the specificity of co-transcriptional pre-miRNA biogenesis and suggests that pre-miR18a and 19a are cleaved post-transcriptionally.

Transcription termination of U snRNA genes is known to be induced by the Integrator complex that recognizes a downstream consensus element (3’Box) and cleaves at adjacent RNA sequence (Chen and Wagner, 2010). Although RNA cleavage sites of transfected or *in vitro* synthesised U snRNA were previously characterised using an RNA protection assay (Uguen and Murphy, 2003), endogenous cleavage positions have not been previously described. Our POINT-5 analysis clearly reveals co-transcriptional RNA cleavage on U snRNA genes by showing S and C peaks, presumably with the C peak derived from Integrator associated endonuclease activity (known to be present in the IntS11 subunit). This is known to cleave pre-U snRNAs immediately upstream of the consensus 3’box sequence (Figures 2D and S2D). It is especially evident for these small Pol II TUs, that 3’processing occurs well before the end of the snRNA primary transcript.

As shown above, strong C peaks were detected for histone gene transcripts, pre-microRNA and U snRNA. This indicates that their 5’ends, even though derived from different co-transcriptional RNA cleavage activities are relatively stable and so detected by POINT-5 technology. Instead we fail to detect C peaks for pre-mRNA PAS. Thus, for *GAPDH*, significant signal was only detected at the TSS (Speak), but not at the TES (C peak) even though endonuclease cleavage at PAS by CPSF-73 is well established (Figure 2E). This led us to a hypothesis that the 5’-3’exonuclease Xrn2 selectively and rapidly degrades downstream RNA following cleavage at the PAS, but not from the other endonuclease cleavage sites. We therefore elected to look further into the mechanism of Xrn2 mediated Pol II termination using POINT-5 technology.

### Specific Xrn2-dependent transcription termination of protein-coding genes

Transcription termination of protein coding genes is well known to be associated with degradation of downstream RNA by the nuclear 5’-3’exonuclease Xrn2, following 3’end CPA of the pre-mRNA (Proudfoot, 2016; West et al., 2004) (Figure 3A). Additional factors cooperate with Xrn2 to promote efficient Pol II termination, both by alteration to the CTD phosphorylation status as well as by the induction of conformational changes to the elongating Pol II complex (Cortazar et al., 2019; Eaton et al., 2020).

**Figure 3.**
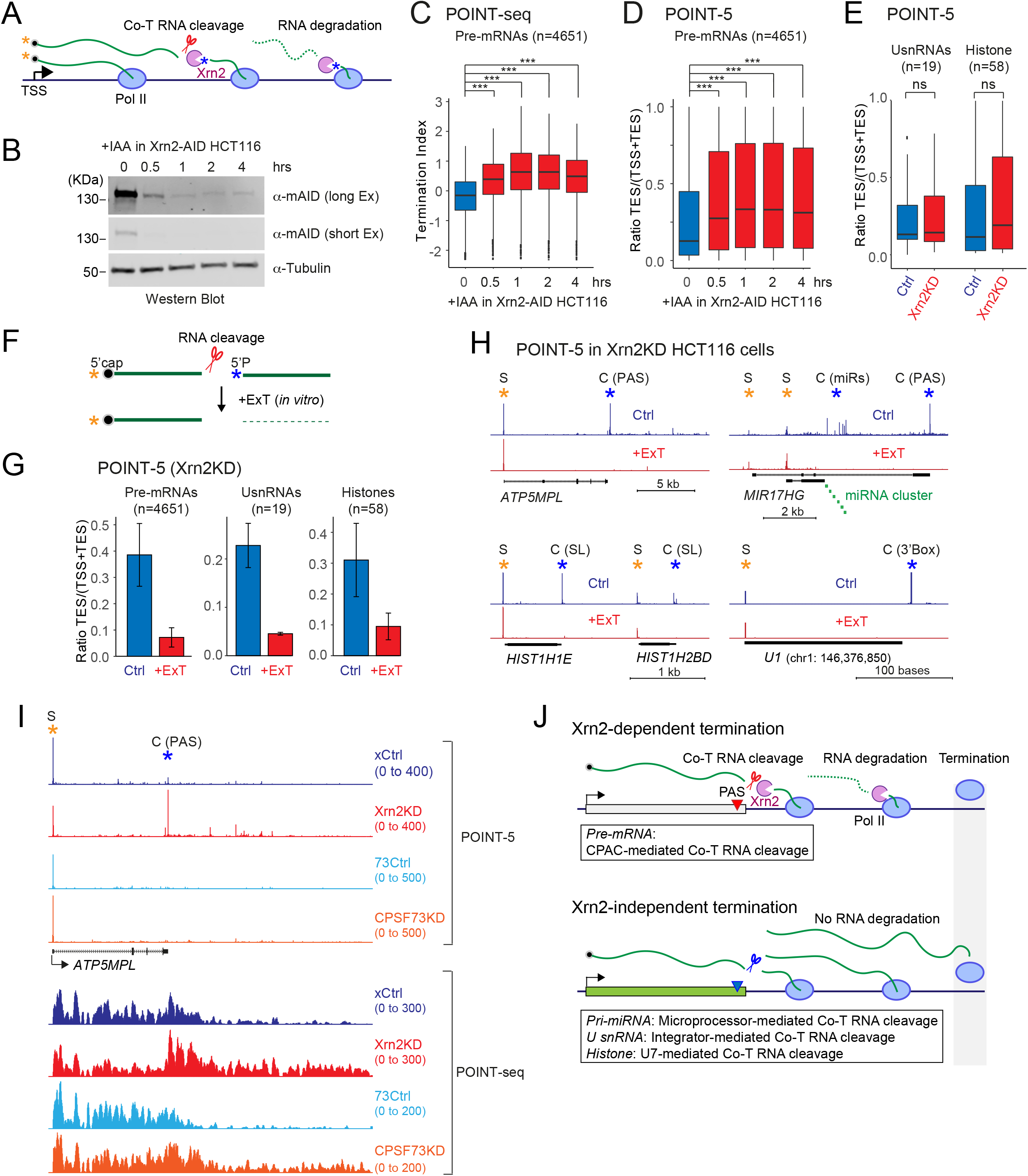
Specificity of Xrn2 exonuclease. (A) Schematic of co-transcriptional RNA cleavage and degradation. Xrn2 recruited at RNA cleavage site and degrades nascent RNA toward Pol II. (B) Western blot of Xrn2-AID over time course of IAA treatment in HCT116 Xrn2-AID cells as indicated (two exposures). Tubulin antibody was used as loading control. (C) Termination Index. Bar plots of POINT-seq signal ratio of gene body and termination region upon Xrn2KD time course for non-overlapping PC genes (n=4,651) expressed in HCT116 Xrn2-AID cells. No treatment (Blue) and Xrn2KD (Red). (D) Bar plots of POINT-5 signal ratio of TSS and TES upon Xrn2KD time course for non-overlapping PC genes (n=4,651) expressed in HCT116 Xrn2-AID cells. No treatment (Blue) and Xrn2KD (Red). (E) Bar plots of POINT-5 signal ratio of TSS and TES upon Xrn2KD 4hr for UsnRNA (n=19) and histone (n=58) genes expressed in HCT116 Xrn2-AID cells. No treatment (Blue) and Xrn2KD (Red). (F) Schematic diagram of ExoTerminator (ExT) treatment. Uncapped 5’P-RNA generated by RNA cleavage and specifically degraded by *in vitro* ExT treatment. (G) Bar plots of POINT-5 signal ratio of TSS and TES upon Xrn2KD 4hr for non-overlapping PC genes (n=4,651), UsnRNA (n=19), and histone (n=58) genes expressed in HCT116 Xrn2-AID cells. No treatment (Blue) and ExT-treated (Red). (H) Examples of POINT-5 upon Xrn2KD (4 hr) with ExT treatment in HCT116 Xrn2-AID cells. (I) *ATP5MPL* as example of POINT-seq and POINT-5 upon Xrn2KD (4 hr) or CPSF73KD (3 hr) in HCT116 Xrn2-AID or CPSF73-AID cells. (J) Schematic model of Xrn2 dependent or independent termination mechanisms.

The role of Xrn2 in processing protein coding gene TES has been previously investigated by use of degron tagged (AID) Xrn2 protein (Eaton et al., 2018). HCT116 cells were engineered to position an AID tag on the endogenous Xrn2 C-terminus as well as to express an ectopic copy of the plant TIR gene that can be activated to degrade Xrn2-AID by auxin treatment. We have employed this Xrn2 degron cell line in our POINT-5 analyses. Note that the AID tag renders Xrn2 unstable even without auxin treatment, so that these cells (Ctrl HCT116) generate substantially less Xrn2 protein that either wild type HCT116 (Eaton et al., 2018) or HeLa cells (Figure S3A). Further, near-complete depletion of Xrn2-AID protein was induced by auxin (IAA) treatment and its levels were monitored over a 4 hr time course by western blot (Figures 3B). Xrn2 protein was substantially depleted following 30 min of IAA treatment. Consistent with previous analysis using mNET-seq (Eaton et al., 2018), POINT-seq shows that rapid depletion of Xrn2 protein induced immediate Pol II termination defects on protein coding genes, with RNA accumulating downstream of PAS (Figures 3C and S3B). Similarly POINT-5 detected an increase in signal at TES regions of protein coding gene following Xrn2 depletion (Figures 3D and S3C). Note that Ctrl HCT116 cells generally show a clear TES peak (Figures S3C and S3D) in contrast to HeLa cells, where TES peaks are barely detected (Figure 1C). This is because expression levels of Xrn2 in HeLa cells are much higher than in Ctrl HCT116 cells caused by the AID tag on Xrn2 inducing partial destabilisation without IAA (Figure S3A). Both defects in transcription termination and reduced RNA degradation at PAS were observed from 30 min IAA treatment (Figures 3C and 3D). This suggests that POINT technology detects direct effects of Xrn2 protein on these transcription events. Xrn2 depletion did not affect the C peaks of either histone and or U snRNA genes, as shown by POINT-5 analysis (Figures 3E and S3E). In addition, POINT-5 C peaks at intronic pre-miRNAs (MIR17HG cluster) were unaffected by Xrn2 depletion, although the C peak of the host gene PAS was significantly increased (Figure S3F). Notably we also detected no effect of Xrn2 depletion on intergenic pre-miRNA (MIR331) even though the pre-miRNA is located downstream of the host gene *VEZT* PAS (Figure S3F). These results underline the fact that Xrn2 specifically degrades nascent RNA cleaved at PAS to induce transcription termination.

Overall the above results indicate that rapid Xrn2 depletion causes termination defects following CPA cleavage of PAS. Since POINT-5 methodology detects all nascent RNA 5’ends irrespective of their chemical nature, we next sought to categorise POINT-5 signals into either 5’capped or uncapped 5’P-RNA. In particular, Xrn2 exonuclease activity is specific for 5’P-RNA substrates. We therefore employed the <u>Ex</u>o<u>T</u>erminator nuclease (ExT) to *in vitro* digest IPed nascent RNA fractions, as isolated by POINT technology. This will selectively degrade Xrn2 sensitive 5’p-RNAs (Figure 3F). Notably POINT-5 TES signals for polyA+ pre-mRNA, U snRNA and histone pre-mRNA were all substantially reduced by *in vitro* ExT treatment, but not TSS-associated S signals (Figures 3G, 3H and S3G). These observations confirm that the POINT-5 TES signals, sensitive to ExT do indeed derive from RNA cleavage and not from alternative TSS. For example, POINT-5 peaks downstream to *JARID2* TSS2 are ExT-sensitive, but not Xrn2-dependent, suggesting that PAS independent RNA cleavage occurs near this alternate TSS of *JARID2* (Figure S3G).

Another feature of rapid Xrn2-AID depletion was the appearance of multiple POINT-5 peaks situated downstream of PAS that all display ExT-sensitivity (Figures 3I and S3G). We hypothesize that these peaks are either intermediates of Xrn2-mediated RNA degradation or RNA cleavage sites of other RNA endonucleases such as Integrator or Drosha that do not induce Xrn2-dependent RNA degradation. We next induced defective transcription termination by CPSF73 degron mediated depletion (Eaton et al., 2020) and determined whether POINT-5 peaks located downstream of PAS are activated by the CPSF73KD. CPSF73 depleted by IAA treatment of HCT116 CPSF73-AID cells for 3 hr (Figure S3H) showed clear termination defects for non-overlapping protein coding genes by POINT-seq analysis (Figure S3I). However, no POINT-5 peaks were detected at PAS even in control cells, because of high Xrn2 protein levels in HCT116 CPSF73-AID cells (Figure S3J). Notably CPSF73 depletion did not induce POINT-5 peaks downstream of *ATP5MPL* PAS (Figure 3I), suggesting that Xrn2-independent RNA degradation does not occur in this termination region. This further indicates that Xrn2 depletion-dependent peaks at the termination site are intermediates of Xrn2-mediated RNA degradation or CPA-cleaved RNA downstream of functional PAS.

Our POINT-technology reveals the precise locations of co-transcriptional RNA cleavage sites and also their Xrn2-dependency (Figure 3J). Protein coding RNA (pre-mRNA) is cleaved by CPA complex at the PAS and then Xrn2 degraded towards elongating Pol II resulting in transcription termination. In contrast, histone pre-mRNA, U snRNA and pri-miRNA are cleaved in a CPA-independent manner. Transcription termination of such genes appears Xrn2-independent, indicating that their 3’end cleaved RNA is undegraded. Possibly these RNA cleavage activities trigger different termination mechanisms such as transcription road block effects caused by loss of transcription elongation factors, DNA structures or higher nucleosome density (Proudfoot, 2016).

### Splicing supresses premature transcriptional termination

Cryptic, inactive PAS are often embedded within mammalian introns. Usage of such intronic PAS may be restricted by mechanisms such as limiting levels of CPA factors or more efficient Pol II elongation (Kamieniarz-Gdula and Proudfoot, 2019). Most importantly, U1 antisense morpholino (AMO) that blocks U1 snRNA base pairing with 5’SS often leads to activation of intronic PAS and consequent premature transcriptional termination (PTT) (Kaida et al., 2010; Oh et al., 2017). The specificity of this effect (referred to as telescripting) was implied by use of U2 AMO which although inhibiting splicing did not induce PTT in a selected gene, normally affected by U1 AMO (Oh et al., 2017). In particular U1 snRNA bound to 5’SS has recently been shown to interact with and inhibit CPA complex activity and so suppress PTT by preventing nearby PAS recognition (So et al., 2019). However, it remains a possibility that splicing may more generally prevent PTT.

We determined the effect of Pladienolide B (PlaB) treatment of HeLa cells on our POINT-seq profiles, as this inhibitor targets the SF3B complex, a component of U2 snRNP. Notably PlaB induced PTT in numerous protein coding genes, but at variable locations across their TUs (Figure 4A). Depending on the positions where POINT-seq signals become interrupted in PlaB treated cells, genes were grouped into four categories: either with PTT located within the first (early, E), second (middle, M) or final (late, L third of each TU. The 4th and dominant category was where no change (NC) in POINT-seq profiles was observed (Figures 4A and 4B). In detail PlaB treatment induced PTT in 21% of non-overlapping protein coding genes expressed in HeLa (Figure 4B) with significant inhibition of splicing in all categories (Figure 4C). Notably PTT induced by PlaB preferentially occurred in long genes (Figure 4D), presumably because Pol II will encounter more cryptic PAS in longer TUs. These results are illustrated for *TBC1D17* (12 kb TU) which shows only splicing inhibition and for *MTHFD1L* (~560kb TU) which shows both splicing and PTT effects following PlaB treatment (Figure 4E).

**Figure 4.**
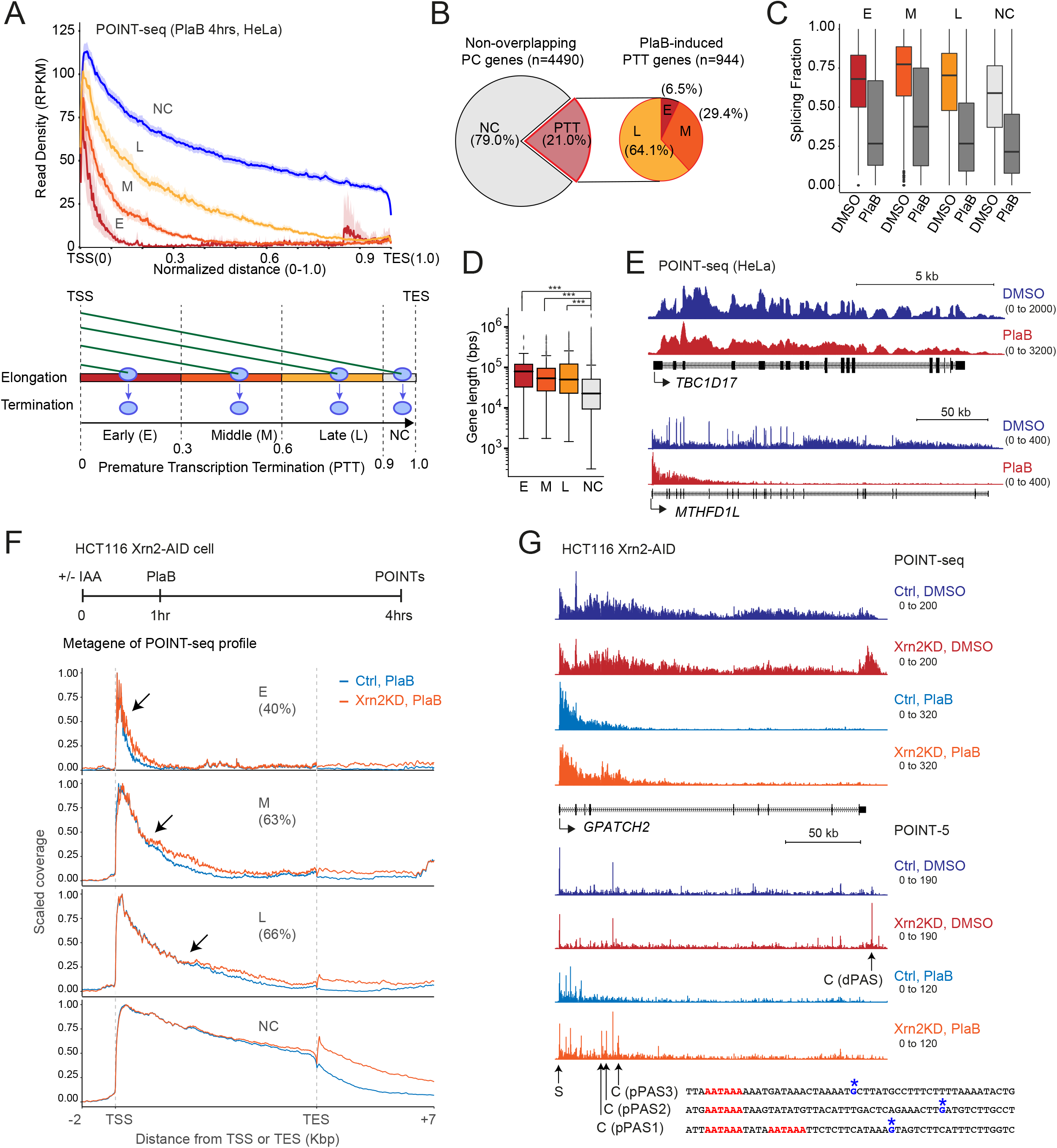
Suppression of PTT by splicing. (A) Four classes (E, M, L, and NC) of premature termination induced by PlaB treatment for 4 hr. (Top) Metagene of POINT-seq in PlaB-treated HeLa cells. (Bottom) Schematic diagram of premature termination in different four regions of normalized gene. (B) Pie charts for fractions of PTT genes induced by PlaB. (C) Splicing fraction in four classes of premature termination genes induced by PlaB. (D) Gene length in four classes of premature termination genes induced by PlaB. (E) Examples of POINT-seq signals on *TBC1D17* (short gene) and *MTHFD1L* (long gene) in HeLa cells. (F) (Top) Schematic protocol of Xrn2KD and PlaB treatment in HCT116 Xrn2-AID cells. (Bottom) Metagene profile of POINT-seq for four classes (E, M, L, and NC) with PlaB (light blue) or “Xrn2KD and PlaB” (orange) in normalized region from TSS-2kbp to TES+7kbp. Percentages of PTT gene cases recovered by Xrn2KD are shown. Arrows indicate start positions of PTT defects induced by Xrn2KD. (G) *GPATCH2* as example of POINT-seq and POINT-5 analyses for indicated treatments. Distal (d)PAS and three cryptic proximal (p)PASs were detected by POINT-5 under Xrn2KD and Xrn2KD + PlaB conditions, respectively.

We next examined whether PlaB induced PTT is caused by splicing inhibition rather than potential indirect effects such as telescripting. We therefore reanalyzed published databases obtained using 4sU metabolically labelled RNA-seq on HeLa cells treated with U1 AMO treatment (8 hr) (Oh et al., 2017). These were compared with our POINT-seq data on HeLa cells treated with PlaB (4 hr). Notably a significant fraction of genes displayed PTT by both U1 AMO and PlaB treatments in HeLa cell (Figure S4A) as shown for *MTHFD1L* where PTT was induced both by U1 AMO and PlaB (Figure S4B). In contrast while *DHX9* shows a clear U1 AMO effect, it does not display PTT effects following PlaB treatment (Figure S4B). The fact that telescripting and general splicing inhibition by PlaB treatment can often affect different genes may indicate different cryptic PAS sensitivities.

We sought to establish that PlaB-induced PTT is regulated by cryptic PAS activation. We therefore depleted Xrn2 or CPSF73 proteins for 1 hr followed by PlaB treatment for 3 hr in the AID engineered HCT116 cells (Eaton et al 2018). POINT-seq analysis of these cells is shown by metagene profiling (Figures 4F and S4C) as well as for the specific gene *GPATCH2* (Figures 4G and S4D). Notably we show that PTT transcripts are indeed extended following depletion of both Xrn2 and CPSF73 proteins. Furthermore, Xrn2 depletion followed by PlaB treatment induced POINT-5 peaks at multiple cryptic PAS (pPAS1-3) in the intron 4 of *GPATCH2* as well as a single POINT-5 peak at the TES (dPAS) in untreated DMSO cells (Figure 4G). We also detected some Xrn2-independent POINT-5 peaks with PlaB treated cells (intron 1 of *GPATCH2*). This implies that CPSF73-independent RNA cleavage and Xrn2-independent RNA degradation may also be involved in premature termination.

### POINT-nano methodology profiles single molecule nascent transcripts

Technical issues have hitherto limited a full understanding of how co-transcriptional RNA processing is regulated during the transcription cycle. Illumina sequencing can be employed to map short nascent RNA fragments, either metabolically labelled by 4sU (Schwalb et al., 2016) or isolated from within Pol II elongation complexes (Nojima et al., 2015). However, read lengths of individual RNA species were generally too short to determine the kinetics of co-transcriptional splicing, interwoven with other RNA processing mechanisms. Indeed, our previous mNET-seq analysis of spliced RNA associated with the Pol II active site was limited to investigation of short exons where immediate upstream splicing was still detected (Nojima et al., 2018a). Here we have employed longer read ONT sequencing on the full-length nascent transcripts that we can isolate using our POINT technology (Figure 5A). Firstly, Pol II-associated intact nascent RNA was isolated using the POINT protocol. RNA longer that 500 nt was size-selected to avoid short reads in ONT sequencing. This size selected RNA fraction was then subjected to *in vitro* 3’end polyadenylation using bacterial derived polyA polymerase. The RNA size range of the *in vitro* pA+ POINT was confirmed by tapestation analysis (Figure S5A). Following reverse transcription with nanopore oligodT primers attached to a motor protein, direct cDNA sequencing was performed using the high throughput ONT device PromethION. Bioinfomatic analysis of POINT-nano required the prior removal of sequence derived from oligodT priming on internal A rich sequence (Figures S5B and S5C) so that only authentic transcript 3’ends, derived from the Pol II active site were considered. We named this strategy for sequencing single molecule, long read transcripts as POINT-nano.

**Figure 5.**
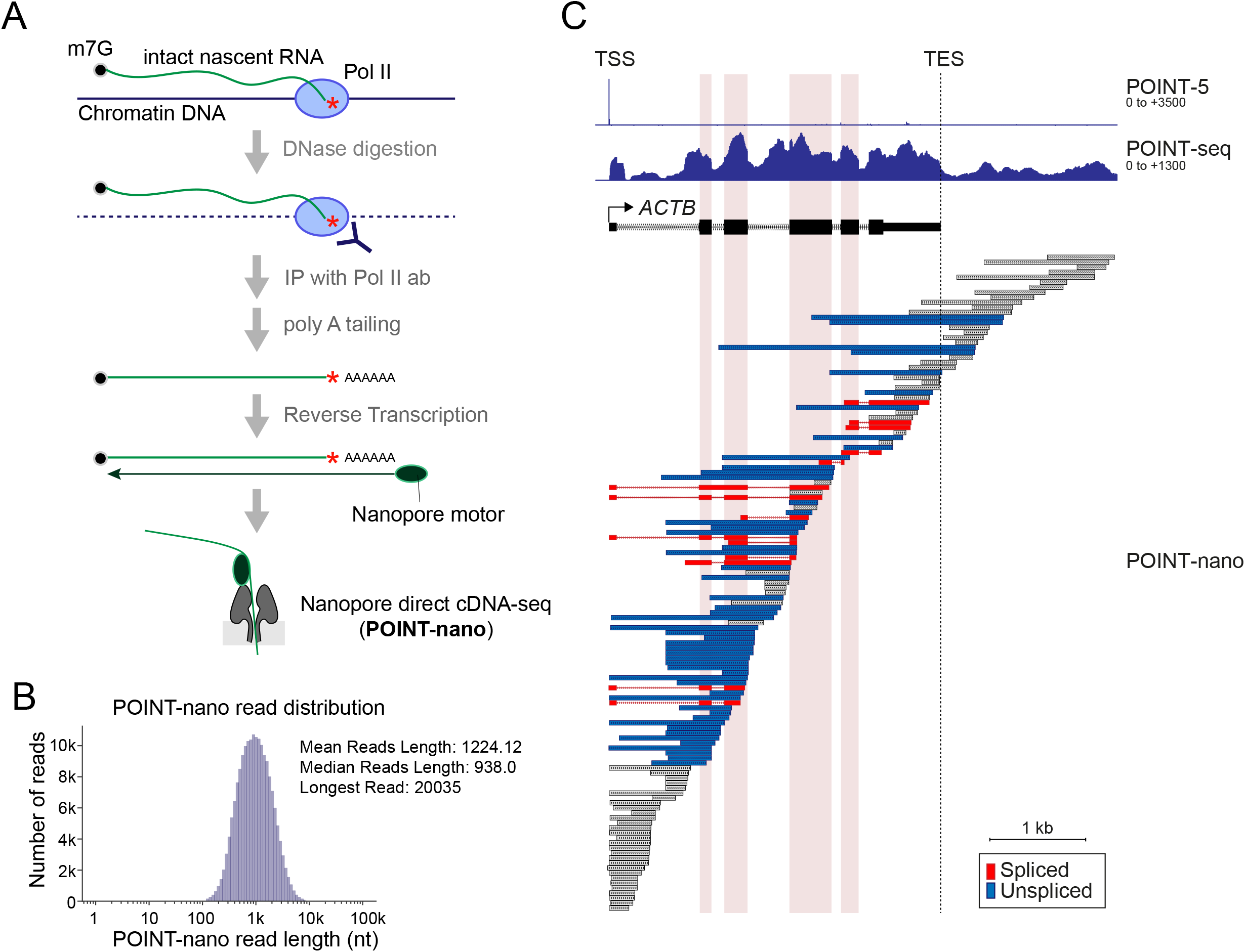
Development of POINT-nano methodology. (A) Schematic of POINT-nano. Intact nascent RNA isolated as in other POINT technologies and *in vitro* polyadenylated. ONT direct cDNA sequencing was applied to poly A tailed intact nascent RNA. (B) Distribution of POINT-nano reads. (C) *ACTB* as example of POINT-nano reads with profiles of POINT-5 and POINT-seq also shown. Internal exons highlighted in light red. TES marked with dashed line. Spliced and unspliced POINT-nano reads indicated in red and blue, respectively. Other POINT-nano reads in grey.

POINT-nano datasets show high reproducibility (Figure S5D) with a mean coverage per non-overlapping protein coding gene of approximately 35 reads (Figure S5E). A comparison of RNA 3’end distributions for POINT-nano versus mNET-seq databases across protein coding gene TUs reveals predominantly intronic locations, which is predicted since introns are much longer than exons in human genes (Figure S5F). Unexpectedly POINT-nano datasets show 5.4% reads at the TES of protein coding genes (Figure S5F). This may indicate that cleaved transcripts remain transiently tethered to elongating Pol II, as also observed in a recent study (Drexler et al., 2020). A restriction to our POINT-nano technology is that read lengths are limited to, on average, 1224 bp (Figure 5B), while the intact nascent RNA have a mean length of ~6,000 nt (Figure S5A).

POINT-nano reads, are shown for *ACTB* and *ID1* and compared to their POINT-5 and POINT-seq profiles (Figures 5C and S5G). These align with PROMPTs and unspliced pre-mRNA, confirming that nascent transcripts are effectively captured by this methodology (Figure S5G). A mix of spliced and unspliced reads are evident on most genes as illustrated for *ACTB* (Figure 5C). Due to the ~1 kb limitation of our POINT-nano data, it is difficult to determine whether read 5’ends correspond to authentic TSS, TES or *in vivo* degradation versus artificial read ends. However multiple reads ending at the TSS as defined by the POINT-5 signal, likely correspond to capped TSS transcript while short reads 3’ to the TES likely correspond to authentic 3’processing by CPA with subsequent Xrn2 degradation. In spite of these read size limitations, our POINT-nano technology can still be employed to profile the status of RNA processing and Pol II position in single RNA molecules, providing new insight into co-transcriptional processing kinetics.

### POINT-nano identifies immediate and delayed co-transcriptional splicing

To determine the timing of pre-mRNA splicing relative to transcription, we grouped spliced and unspliced reads detected by POINT-nano sequencing according to the location of their 3’ ends in either exons or introns (Figure 6A). The results show that substantial fraction of nascent RNA molecules are already spliced as Pol II transcribes an exon downstream of the 3’SS (Figure 6B), indicative of splicing by intron definition. As expected, the proportion of spliced POINT-nano reads was greatly reduced by PlaB treatment, based on bioinformatic measurement of the splicing fraction (Figure 6B) or by focusing on products of *SRSF2* transcription (Figure 6C). Next, we analyzed the proportion of nascent transcripts that become spliced as Pol II transcribes across the 3’ and 5’SSs in Ctrl DMSO and PlaB-treated cells. To circumvent bias introduced by read length, we restricted the analysis to a fixed number of nucleotides around each SS. Compared to intronic regions, a clear enrichment of spliced reads was observed on exons (Figure 6D). Notably, some spliced reads were detected as early as 15nt downstream of 3’SS (Figure S6A), which is consistent with the number of newly synthesized nucleotides located within the Pol II complex (Martinez-Rucobo et al., 2015). This suggests that splicing can occur immediately after the nascent transcript emerges from the RNA exit channel of Pol II. We further observed that the proportion of spliced reads decreased abruptly at the exon-intron boundary, but then increased as Pol II transcribed along the downstream intron (Figures 6D and S6B). These results indicate that POINT-nano detected two different types of spliced RNA molecules. We therefore conclude that splicing of newly synthesized pre-mRNA can occur either immediately after transcription of the 3’SS (while Pol II is located on the exon) or be delayed until Pol II transcribes several thousand base pairs of the downstream intron (Figure 6E), in agreement with the view that splicing can be delayed until exon definition occurs.

**Figure 6.**
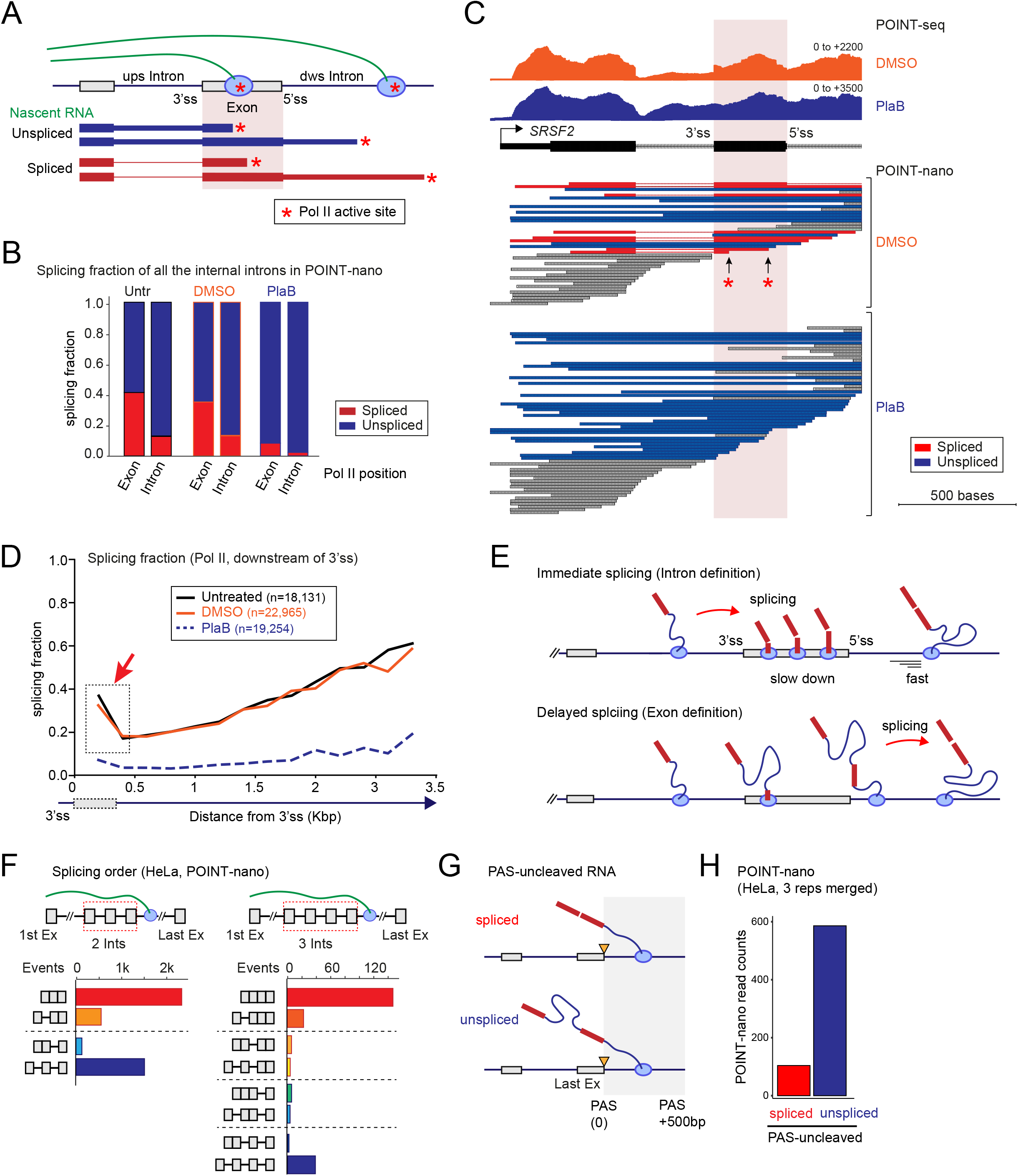
Two modes of co-transcriptional splicing in human cells. (A) Schematic of POINT-nano analysis on timing of co-transcriptional splicing. Splicing of downstream exon highlighted in light red. Spliced and unspliced POINT-nano reads indicated in red and blue, respectively. Pol II active site indicated by red asterisk. (B) Splicing fraction of POINT-nano reads for all internal introns expressed in untreated (Untr), DMSO and PlaB-treated HeLa cells. Pol II positions indicated below. (C) *SRSF2* as example of POINT-nano reads with profiles of POINT-seq in DMSO and PlaB-treated HeLa cells. Spliced and unspliced POINT-nano reads indicated in red and blue, respectively. Other POINT-nano reads in grey. Pol II active sites on spliced exon 2 indicated as red asterisks. (D) Splicing fraction of POINT-nano reads with Pol II located downstream of 3’SS. Untreated, DMSO and PlaB-treated HeLa cell shown as black, orange and blue dashed lines. High splicing fraction near 3’SS indicated in dashed line box with red arrow. (E) Two co-transcriptional splicing models, immediate and delayed splicing in human. (F) Splicing order analysis of POINT-nano reads for (left) two and (right) three internal introns in HeLa cell. (G) Diagram illustrating spliced/unspliced PAS-uncleaved transcripts over TES region. (H) Splicing fraction for last intron removal in POINT-nano analysis. Pol II positions indicated below.

We further investigated the ability of POINT technology to discriminate differences in splicing efficiency associated with distinct SSs. Notably, the proportion of spliced reads aligned with weak SSs was significantly lower compared to reads spanning strong SSs (Figures S6C and S6D). Since alternatively spliced exons tend to have weaker SSs (Itoh et al., 2004), we determined whether POINT technology detects differences in co-transcriptional splicing efficiency between constitutive and alternative splicing events. For this analysis we employed POINT-seq data since POINT-nano datasets had a very low read coverage. The results show that the proportion of spliced reads is significantly higher in constitutively spliced exons compared to alternative exons (Figure S6E). In order to determine the timing of alternative splicing, we focused on cassette exon splicing events. These were classified into high or low exon inclusion categories using previously published pA+ RNA-seq data sets (Figure S6F). As expected, our POINT-nano data reiterated this classification. Importantly, whether or not the internal cassette exon was included or excluded, splicing levels of the external exons were maintained (Figure S6G). This suggests that both exon inclusion and exclusion events can be regulated by immediate splicing (i.e. intron definition). Notably we cannot exclude the possibility that cassette exon splicing can also by regulated by delayed splicing (i.e. exon definition) since our POINT-nano data does not detect RNAs longer than 10,000 nt (Figure 5B) due to technical limitations as described above.

We next used POINT-nano to investigate splicing order along the TU. Due to the limited average length of POINT-nano reads, we analyzed the order of excision of either two or three internal introns (Figure 6F). In the majority of cases all introns were excised (Figure 6F, red bars). However, sometimes an upstream intron was left unspliced even though the downstream intron was excised (Figure 6F, orange bars). This demonstrates that splicing does not always occur sequentially. We further detected a major subset of transcripts with all analyzed introns remaining unspliced (Figure 6F, blue bars). To investigate the timing of splicing relative to 3’end cleavage and polyadenylation, we analyzed POINT-nano reads with 3’ends located within 500 bp past the TES (Figure 6G). We detected multiple reads corresponding to nascent transcripts not cleaved at the PAS, and notably the vast majority of these were unspliced (Figure 6H). Consistent with this, we also detected reduced splicing of the last intron when Pol II is located downstream of PAS compared to when it is in the last exon (Figure S6H). Whether such transcripts will eventually be processed and exported to the cytoplasm or targeted for degradation in the nucleus remains to be established.

### Extending TU size enhances co-transcriptional splicing

We observe that the proportion of spliced reads gradually increase as Pol II transcribes further into the downstream intron (Figure 6D) and so hypothesized that intron length enhances splicing level. To better understand this, we classified exons and introns according to their size into either long (top 25%) or short (bottom 25%) and estimated the proportion of spliced POINT-nano reads in these two categories. Notably splicing level was similar for both short and long exons, even though the distance from the 3’SS at which spliced reads decline was predictably shorter for the shortest exons (Figure S7A). Similar levels of spliced reads were detected on the downstream intron irrespective of exon size (Figure S7B). In contrast, longer introns were associated with a higher proportion of spliced reads corresponding to both immediate (Figure S7C) and delayed splicing (Figure S7D). This effect is statistically significant (Figure S7E). Since long introns are generally present in long genes, we reasoned that long genes should be more efficiently spliced than short genes. POINT-seq datasets confirm that indeed the proportion of spliced reads increased with gene length (Figure 7A). As a control, we calculated the proportion of spliced reads detected in libraries prepared from pA+ RNA fraction which gave largely spliced reads irrespective of gene length (Figure 7A).

**Figure 7.**
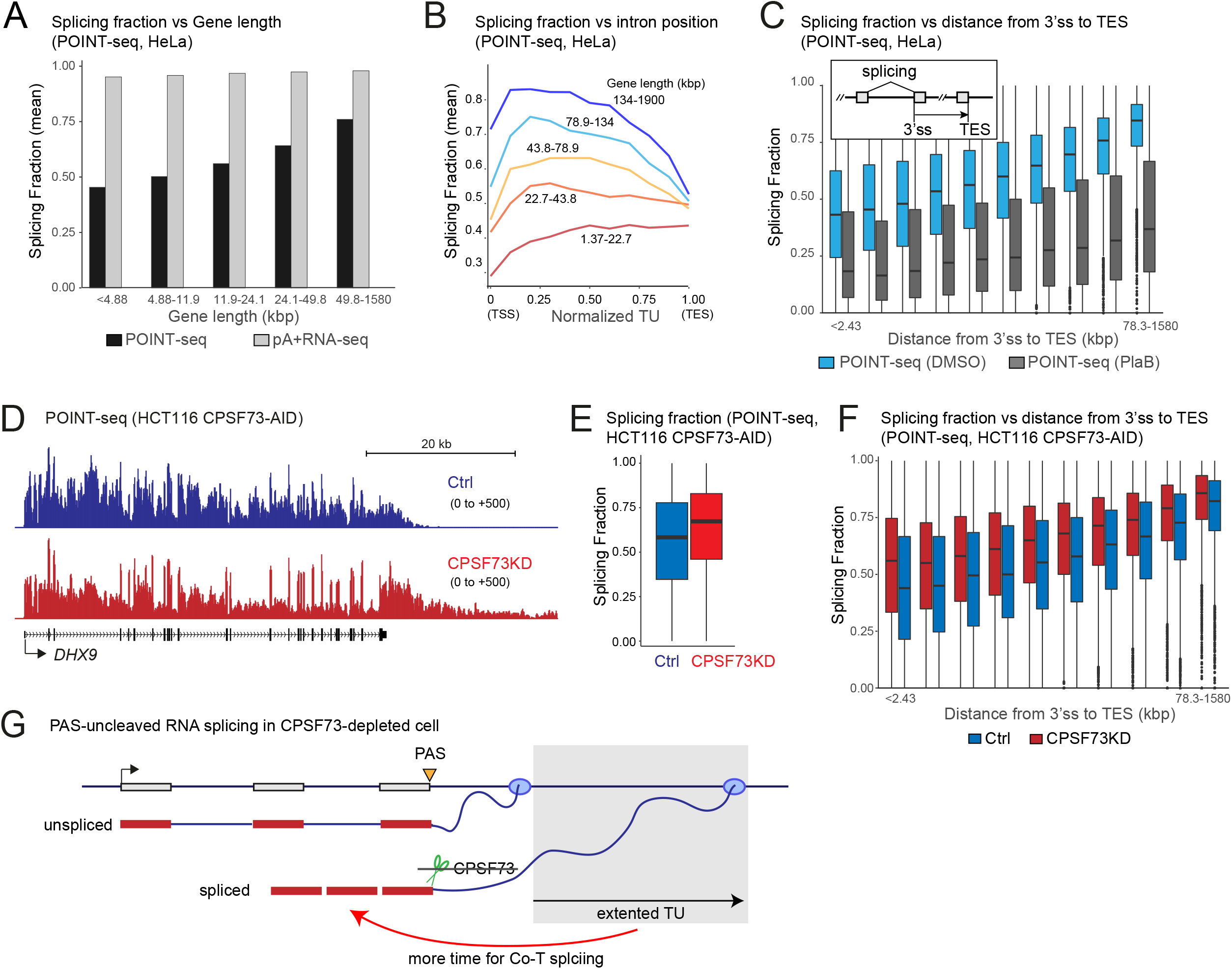
Co-transcriptional splicing following RNA cleavage at PAS. (A) Splicing fraction of POINT-seq (black) and pA+ RNA-seq (grey) from HeLa cells with different gene lengths as indicated. (B) Splicing fraction in each intron positions of normalized TU with different gene lengths as indicated. POINT-seq analysis in HeLa cells. (C) Splicing fraction of POINT-seq from DMSO (light blue) or PlaB (dark grey)-treated HeLa cells with different distance from 3’ss to TES as indicated. (D) *DHX9* as example of POINT-seq profiles upon CPSF73KD in HCT116 CPSF73-AID cell. (E) Splicing fraction of POINT-seq signals upon CPSF73KD in HCT116 CPSF73-AID cell. (F) Splicing fraction of POINT-seq signals upon CPSF73KD in HCT116 CPSF73-AID cell with different distance from 3’SS to TES as indicated. (G) Schematic model of co-transcriptional (Co-T) splicing regulated by defects of RNA cleavage and transcription termination. Orange triangle is PAS. Green scissor shows RNA cleavage activity of CPSF73 protein. Red arrow head shows enhancement of Co-T splicing.

We next analyzed the proportion of spliced reads associated with introns located at different positions along the TU (Figure 7B). We found that splicing fraction of the first intron is generally lower than internal introns as previously reported in *S. pombe* (Herzel et al., 2018). Moreover, splicing fraction gradually decreased from the middle of the TU towards the TES, except for the shortest gene class (Figure 7B). This suggests that splicing level depends on the time that Pol II spends elongating past the 3’SS. Indeed, analysis of POINT-seq datasets shows that the proportion of spliced reads increased with the distance from 3’SS to TES (Figure 7C). PlaB treatment confirmed the authenticity of spliced reads detected by POINT-seq. We also analyzed the effect on splicing of the distance to TES from 3’SSs located at different positions along a normalized TU. This shows similar effects for all positions with both constitutive and alternatively spliced exons (Figures S7F and S7G).

Our bioinformatic analysis of splicing shows a clear kinetic effect whereby longer introns or longer genes allow more time to complete successful splicing. We therefore reasoned that inducing an artificial extension to Pol II TUs should also act to enhance splicing level. Consequently we depleted CPSF73 to prevent normal TES recognition and measured spliced read levels by POINT-seq. Degradation of CPSF73 was induced in HCT116 CPSF73-AID cells by addition of IAA for 3 hr (Fig. S3H) and our POINT-seq analysis confirmed that this caused read-through transcription at gene ends as shown for *DHX9* (Figure 7D). We further observed that upon CPSF73 depletion, transcription termination defects at the end of *HNRNPA0* caused transcript “read-in” on normally silent *KLHL3* (Figure S7H). Interestingly spliced reads corresponding to *KLHL3* transcripts were detected, indicating that Pol II termination defects do not inhibit splicing. We then focused on the proportion of spliced reads along protein coding genes and found a significant increase in CPSF73 depleted cells (Figures 7E and S7I). Enhanced splicing level was observed throughout internal splicing events along the TU, as shown by a relative enrichment of exonic signals in *DHX9* transcripts (Figure 7D) and by quantitative analysis of the proportion of spliced reads at different distances from the TES (Figure 7F). Overall our analysis of extended TUs caused by CPSF73 depletion reveals that upstream splicing efficiency is enhanced presumably by providing more time for assembly of a catalytically active spliceosome (Figure 7G).

## DISCUSSION

### POINT methodology

Several sequencing technologies have been developed to profile mammalian nascent transcription (Stark et al., 2019). GRO-seq and PRO-seq employ *in vitro* nuclear run on (NRO) transcription with modified ribonucleotides to label and isolate newly synthesized RNA (Core and Lis, 2008; Kwak et al., 2013). Metabolic labelling of nascent RNA can also be carried out on cell cultures by application of 4sU-seq and TT-seq (Schwalb et al., 2016; Windhager et al., 2012). These methods have been widely employed to dissect Pol II pausing, elongation speed, and RNA stability. Furthermore, sequencing chromatin associated RNA 3’ends can also be employed as a surrogate to sequence Pol II active site associated transcripts (Mayer et al., 2015). However, it is apparent that chromatin associated RNA contains significant amounts of contaminating steady state mRNA (Schlakow et al 2017). We previously developed mammalian mNET-seq (Nojima et al., 2015) which effectively circumvents nucleoplasmic contamination issues. In mNET-seq, both accessible DNA and RNA in the chromatin-fraction is digested by MNase treatment and the transcription machinery is then IPed with Pol II antibody from the solubilized chromatin. Sequencing the protected short RNA fragment within the Pol II complex effectively defines the nascent RNA 3’end. mNET-seq has allowed the characterisation of Pol II pausing and the detection of splicing intermediates (Nojima et al., 2015; Nojima et al., 2018a). However, the restricted read length obtained by mNET-seq has precluded the analysis of RNA processing kinetics in mammalian cells. Here we describe POINT technology which provides a new method to profile intact and truly nascent RNA. Notably a rRNA depletion step routinely employed in most RNA-seq methodologies is unnecessary for POINT technology. This is because a high concentration of strong detergent (3% Empigen) is added during both chromatin isolation and Pol II IP steps which removes all contaminating RNA species without affecting Pol II IP. These contaminated RNA such as mature spliced transcripts are often detected in other nascent RNA labeling approaches (Figure S4B). Consequently, POINT technology enables the rapid purification of a truly nascent RNA preparation. As a way to emphasize the utility of our POINT technology, we have directly employed POINT-5, POINT-seq and POINT-nano to better understand how co-transcriptional RNA processing of nascent transcription by RNA polymerase II is executed.

### POINT-5 analysis reveals different categories of transcript cleavage

POINT-5 provides powerful methodology to map transcript 5’ends that rivals the widely employed CAGE technique (Figures 1E and 1F). CAGE has been applied to map TSS of steady state RNA and in some cases chromatin-bound RNA (Hirabayashi et al., 2019). Notably our POINT-5 method not only defines TSS positions, but also detects co-transcriptional RNA cleavage sites corresponding to 5’p-RNA. Furthermore, we can distinguish RNA cleavage sites from TSS by use of the 5’P specific exonuclease ExT. We show that RNA cleavage at PAS is only readily detected by POINT-5 following Xrn2 depletion. This confirms that the downstream product of PAS cleavage is rapidly degraded by Xrn2 to promote Pol II transcriptional termination (Proudfoot, 2016). Instead POINT-5 analysis detected 3’ end cleavage of pri-miRNA, pre-U snRNA, and pre-histone transcripts without prior Xrn2 depletion. For these gene classes, it is known that RNA cleavage is required to induce their transcription termination. However Xrn2 loss did not significantly affect transcription termination of U snRNA and histone genes in POINT-seq (Figure S3E) as previously shown by mNET-seq analysis (Eaton et al., 2018). Either these cleaved RNAs are not subject to degradation or possibly are degraded by an unknown 5’-3’exonuclease. A possible exonuclease candidate is Xrn1 which although mainly localized in the cytoplasm, may also possess nuclear functions (Sohrabi-Jahromi et al., 2019).

PTT is a critically important step to fine tune gene expression (Kamieniarz-Gdula and Proudfoot, 2019). Previously U1 snRNP was shown to suppress cleavage at intronic cryptic PAS and thereby restrict PTT by a mechanism referred to as telescripting (Kaida et al., 2010). However, we find that a significant number of genes with PTT suppressed by U1 snRNP are also regulated by SF3B1, a component of U2 snRNP (Figure S4A). This suggests that splicing is often directly connected to PTT. Additionally, such PTT induced by PlaB is significantly extended by CPSF73KD (55% of gene cases) and Xrn2KD (65% of gene cases), suggesting that in these cases PTT is PAS-dependent. In contrast we also observe Xrn2-independent POINT-5 peaks, sensitive to ExT treatment near the TSS of protein coding genes (Figure S3G). Possibly the Integrator complex is responsible for these cleavage sites as a way to regulate PTT. Indeed recent studies have shown that Integrator complex prematurely terminates protein coding genes (Elrod et al., 2019; Tatomer et al., 2019) and also terminates transcription of long noncoding RNA genes (Lai et al., 2015). This suggests that the PTT may be regulated by a combination of CPA and Integrator complexes, similar to termination of lncRNA transcription (Nojima et al., 2018b).

### POINT-nano defines different kinetic classes of co-transcriptional splicing

Based on our POINT-nano analysis, we observe two main categories of co-transcriptional splicing; immediate and delayed. Firstly splicing can occur to a 40% level immediately following intron synthesis, with elongating Pol II still within the downstream exon. Instead if elongating Pol II reaches the following intron, then splicing levels are initially lower, but gradually increase as Pol II elongates through the intron and further into 3’regions of the TU. Notably high-resolution ChIP-seq and mNET-seq data reveals that Pol II CTD phosphorylated on serine 5 positions (S5P) is enriched over exons where it acts to recruit active spliceosomes. Subsequently, CTD phosphorylation status changes to S2P as Pol II elongates into downstream introns (Chathoth et al., 2014; Nojima et al., 2018a). Such a CTD phosphorylation transition may be important to control Pol II elongation speed during transcription. Indeed, our current model is that Pol II is paused over exons to allow immediate splicing but then speeds up following completion of splicing, likely due to the release of the spliceosome from the elongation complex. Nucleosome positioning may also contribute to regulation of immediate splicing by slowing down Pol II in intron-exon boundary (Schwartz et al., 2009; Tilgner et al., 2009). We note that a related study using ONT did not detect immediate splicing in human cells (Drexler et al., 2020). However, their use of 4sU-labeling to isolate nascent RNAs could potentially suppress splicing. Also, their chromatin derived RNA may be significantly contaminated with rRNA and mature mRNA as described above and so may underscore the levels of co-transcriptional splicing.

Another notable feature of our splicing analysis by POINT-nano is that splicing increases to higher levels with longer TUs. Thus, either greater intron or gene size correlates with increased levels of splicing. These correlations imply a simple kinetic model whereby more Pol II elongation time gives more time for productive spliceosome assembly or splicing completion on upstream introns. However, we also note that a fraction of transcripts appears refractory to both splicing and PAS mediated 3’end processing. We predict such transcripts may be defective and likely subjected to degradation by nuclear quality control. As a way to test the hypothesis that longer TUs correlate with higher splicing efficiency we tested the effect of abrogating PAS cleavage by CPSF73 depletion. This data confirmed that extending TU length by preventing Pol II termination did indeed promote more efficient splicing of upstream introns. This albeit artificial situation may have relevance to the regulation of gene expression though alternative PAS selection. In particular a switch from proximal to distal PAS that is correlated with cellular differentiation (Di Giammartino et al., 2011) may be able to stimulate upstream splicing events.

Overall, we present POINT technology as a powerful approach to dissect the complex life of nascent RNA, its synthesis, RNA processing and degradation. The POINT-5 method provides a valuable approach to profile co-transcriptional RNA cleavage at single nucleotide resolution. Furthermore, POINT-nano has high potential to reveal more detailed interplay between co-transcriptional splicing and RNA 3’end processing. An improvement in ONT sequencing platforms to afford greater read length and genomic coverage for direct cDNA and RNA analysis remains critical. Even so we anticipate that our POINT technology has high potential to achieve a much deeper understanding of the mechanisms and kinetics of coupled transcription and RNA processing. The human population is increasingly afflicted by pandemics induced by viruses such as HIV, influenza and coronavirus. We anticipate that POINT technology will provide a new tool to understand complex co-transcriptional events in newly emerging human pathogen and their host cells.

## STAR METHODS

Detailed methods are provided in the online version of this paper as following.
- KEY RESOURCES TABLE
- CONTACT FOR REAGENT AND RESOURCE SHARING
- EXPERIMENTAL MODEL AND SUBJECT DETAILS
- METHOD DETAILS

○ AUXIN-DEPENDENT PROTEIN DEPLETION
○ POINT METHODS AND LIBRARY PREPARATION
- QUANTIFICATION AND STATISTICAL ANALYSIS

○ ILLUMINA DATA PRE-PROCESSING
○ POINT-NANO PRE-PROCESSING
○ IDENTIFICATION OF EXPRESSED GENES
○ METAGENE ANALYSIS
○ HEATMAPS
○ CLEAVAGE RATIO
○ TERMINATION INDEX
○ PREMATURE TRANSCRIPTION TERMIANTION ANALYSIS
○ SPLICING ANALYSES WITH POINT TECHNOLOGY
○ ALTERNATIVE SPLICING AND CASSETTE CASES INDENTIFICATION
○ SIGNAL EXTRACTION AND DATA VISUALIZATION
○ READS QUANTIFICATION
○ P-VALUES AND SIGNIFICANCE TESTS

## SUPPLEMENTAL INFORMATION

Supplemental information can be found online.

## ACKNOWLEDGMENTS

We thank members of the NJP and MC-F groups, especially Dr. Rosina Savisaar for critical discussion. We thank Dr. Steven West for providing us HCT116 Xrn2-AID and CPSF73-AID cells and critical comments on this paper. We also thank Dr. Carika Weldon for helping ONT sequencing. This work was supported by funding to NJP (Wellcome Trust Investigator Award [107928/Z/15/Z] and ERC Advanced [339270] Grants) and to MC-F and RS-L (Fundação para a Ciência e a Tecnologia, Portugal, Grant PTDC/MED-ONC/29469/2017 and Fellowship SFRH/BD/147906/2019, respectively). Funding was also received from the European Union’s Horizon 2020 Research and Innovation Programme (RiboMed, 857119).

## AUTHOR CONTRIBUTIONS

RS-L conducted all bioinformatic analyses. TN performed all molecular biology experiments, with help from GD for biochemical experiments and interpretation of splicing data. IZ performed Xrn2 western blot. HK provided valuable Pol II antibodies. TN, NJP and MC-F designed this project. RS-L, NJP, MC-F, and TN wrote the paper.

