## Supplemental info for "POINT Technology Illuminates the Processing of Polymerase-Associated Intact Nascent Transcripts"

#### STAR METHODS

#### KEY RESOURCES TABLE

| REAGENT OR RESOURCE | SOURCE | IDENTIFIER |
|---|---|---|
| Reagent | | |
| Auxin (IAA) | Sigma | Cat# I2886 |
| Tetracycline (Tet) | Sigma | Cat# 87128 |
| Empigen ~30% | Sigma | Cat# 30326 |
| Turbo DNase | Thermo Fisher Scientific | Cat# AM2239 |
| RiboLock RNase inhibitor | Thermo Fisher Scientific | Cat# EO0381 |
| Dynabeads M280 Sheep Anti-mouse IgG | Thermo Fisher Scientific | Cat# 11202D |
| *E. coli* Poly A polymerase | NEB | Cat# M0276 |
| Pladienolide B | Santa Cruz | Cat# SC-391691 |
| DRB | Sigma | Cat#D1916 |
| IAA | Sigma | Cat#I2886 |
| SPRISelect reagent | Beckman Coulter | Cat# B23317 |
| Spike-in SIRV-Set2 | Lexogen | Cat#050.0 |
| Antibodies | | |
| Mouse monoclonal anti-Pol II CTD, Total | (Nojima et al., 2015) | CMA601, Available from Kimura Lab by request. |
| Mouse monoclonal anti-H3 | Active Motif | Cat# 39763 |
| Xrn2 | Bethyl Laboratories | Cat# A301-101A |
| Tubulin | Abcam | Cat# ab7921 |
| mAID | MBL | Cat# M214-3 |
| Deposited Data | | |
| Raw sequencing data | This paper | GEO: GSE159326 |
| Re-analysed ChrRNA-seq data | (Nojima et al., 2015) | GEO: GSE60358 |
| Re-analysed pA+ RNA-seq HeLa S3 data | http://www.genome.gov/27528022 | GEO: GSE86661 |

| | | |
|---|---|---|
| Re-analyzed pA+ RNA-seq HCT116 data | http://www.genome.gov/27528022 | GEO: GSE33480 |
| Re-analyzed U1 AMO 4shU-seq data | (Oh et al., 2017) | GEO: GSE103252 |
| Re-analyzed CAGE data | (Chen et al., 2016) | GEO: GSE75183 |
| Cell Lines | | |
| HeLa (human) | Proudfoot Lab | N/A |
| Xrn2-AID HCT116 (human) | (Eaton et al., 2018) | Available from West Lab by request. |
| CPSF73-AID HCT116 (human) | (Eaton et al., 2020) | Available from West Lab by request. |
| Gels | | |
| Novex 6% TBE gel, 12 well | Invitrogen | Cat# EC62652BOX |
| 4-15% Mini-PROTEAN Precast Protein Gels, 12 well | BioRad | Cat# 4561085 |
| Kits | | |
| NEBNext Ultra II Directional RNA library prep kit for illumina | NEB | Cat# E7760S |
| SMARTer Stranded RNA-seq kit | Takara Bio | Cat# 634836 |
| Direct cDNA library prep kit | Oxford Nanopore Technologies | Cat#SQK-DCS109 |
| PromethION48 flow cell | Oxford Nanopore Technologies | Cat# FLO-POR002 |
| Nanopore barcoding kit | Oxford Nanopore Technologies | Cat# EXP-NBD114 |
| Direct-zol RNA microprep | Zymo Reseach | Cat# R2061 |
| Software and Algorithms | | |
| FastQC (v0.11.5) | | https://www.bioinformatics.babraham.ac.uk/projects/fastqc/ |
| TrimGalore (v0.4.4) | | https://www.bioinformatics.babraham.ac.uk/projects/trim_galore/ |

| | | |
|---|---|---|
| STAR (v2.7.0) | | https://github.com/alexdobin/STAR |
| SAMtools (v1.9) | | http://www.htslib.org/ |
| BedTools (v2.29.2) | | http://bedtools.readthedocs.io/en/latest/content/installation.html |
| Deeptools (v3.4.3) | | http://deeptools.readthedocs.io/en/latest/index.html |
| Guppy Basecalling (v3.0.5) | Oxford Nanopore Technologies | -- |
| NanoQC (v0.9.1) | | https://github.com/wdecoster/nanoQC |
| qcat (v1.1.0) | Oxford Nanopore Technologies | https://github.com/nanoporetech/qcat |
| regEx (v2.5.76) | | https://pypi.org/project/regex/ |
| Porechop (v0.2.4) | | https://github.com/rrwick/Porechop |
| minimap2 (v2.17-r941) | | https://github.com/lh3/minimap2 |
| pysam (v0.15.4) | | https://github.com/pysam-developers/pysam |
| Kallisto (v0.46.0) | | https://github.com/pachterlab/kallisto |
| ggplot (v3.3.2) | | https://cran.r-project.org/web/packages/ggplot2/index.html |
| MaxEntScan | | http://hollywood.mit.edu/burgelab/maxent/Xmaxentscan_scoreseq.html |
| scales (v1.1.1) | | https://scales.r-lib.org/ |
| vast-tools (v2.5.1) | | https://github.com/vastgroup/vast-tools |
| bedGraphToBigWig (v4) | | https://www.encodeproject.org/software/bedgraphtobigwig/ |

#### CONTACT FOR REAGENT AND RESOURCE SHARING

Further information and requests for resources and reagents should be directed to Takayuki Nojima.

### EXPERIMENTAL MODEL AND SUBJECT DETAILS

HeLa and HCT116 cells were maintained in high glucose Dulbecco's Modified Eagle's Medium (DMEM) with 10% fetal bovine serum (FBS) and penicillin/streptomycin (PS).

### METHOD DETAILS

##### Auxin-dependent protein depletion

IAA (final concentration 0.5 mM) was directly added to Xrn2-AID HCT116 cells in DMEM/10%FBS/PS and incubated for 30 min-4 hr as previously published (Eaton et al., 2018). For CPSF73 protein depletion, CPSF73-AID HCT116 cells were incubated with Tetracycline (final concentration 1 μg/mL) in DMEM/10%FBS for 18 hr and then IAA was treated for 3 hr as previously published (Eaton et al., 2020).

##### POINT methods and library prep

POINT method was performed as previously described mNET-seq protocol (Nojima et al., 2016) with some alterations. In brief, crude nuclear fraction was prepared from HeLa or HCT116 cells ($1x10^7$ for POINT-seq and POINT-5, $4x10^7$ for POINT-nano) as in mNET protocol. Chromatin pellet was resuspended in NUN1 (20 mM Tris-HCl (pH 7.9), 75 mM NaCl, 0.5 mM EDTA and 50% Glycerol) and treated with modified NUN2 buffer (20 mM HEPES-KOH (pH 7.6), 300 mM NaCl, 0.2 mM EDTA, 7.5 mM $MgCl_2$, 1% NP-40, 1 M Urea, 3% Empigen, 1x proteasome inhibitor Complete (-EDTA) and 1x PhosSTOP). Note Empigen-treated nuclear solution was gently mixed by inverting tube a few times to avoid chromatin aggregation and incubated on ice for 10 min. The chromatin pellet was centrifuged at 400g for 30 sec. The pellet was washed with PBS twice and then digested in DNase solution (10 mM Tris-HCl (pH 7.5), 400 mM NaCl, 100 mM $MnCl_2$, 2 U/μL RiboLock and 0.2 U/μL Turbo DNase) for 15 min. After DNA digestion, soluble digested chromatin was collected by 13,000 rpm centrifugation for 10 min. The supernatant was ten times-diluted in ice-cold NET-2E buffer (50 mM Tris-HCl (pH 7.4), 150 mM NaCl, 0.05 % NP-40 and 3% Empigen BB) and Pol II antibody-conjugated beads were added. 10 or 40 μg of Pol II antibody (200 or 800 μL of Dynabeads anti-mouse IgG) was used for POINT-seq and POINT-5 or POINT-nano methods. Immunoprecipitation was performed at 4°C for 1 hr. The beads were washed with 1 ml of ice-cold NET-2E buffer six times. The isolated nascent RNA was purified using Trizol reagent technology twice with once DNase treatment. Note that polyA tails were added to the isolated RNA by *in vitro* polyadenylation with *E.coli* PAP for POINT-nano. Then the pA+RNA was size-selected using SPRISelect reagent (x0.6 volume). Library prep for illumina sequencing (NovaSeq6000) employed NEBNext Ultra II Directional RNA library prep kit and SMARTer Stranded RNA-seq kit for POINT-seq and POINT-5 analyses, respectively. For ONT sequencing, Direct cDNA library prep kit was used for POINT-nano analysis. Illumina and ONT sequencing (PromethION) were conducted by Novogene UK and the high throughput genomics team of the Wellcome Trust Centre for Human Genetics (WTCHG), Oxford.

#### QUANTIFICATION AND STATISTICAL ANALYSIS

##### Illumina data pre-processing

Quality control for raw short-reads was performed over POINT-seq and POINT-5 data using the FastQC tool. Then, read adaptors were trimmed using TrimGalore in paired-end mode, removing reads with less than 10 bases and/or low-quality ends (20 Phred score cut-off). The resultant reads were aligned against the reference human genome (GRCh38) using STAR software (Dobin et al., 2013), requiring uniquely mapped reads (--outFilterMultimapNmax 1) and minimum alignment score (--outFilterScoreMin) of 10. Additionally, for POINT-5 the 5'end of the original RNA and their directionality was extracted. To do this, the script created for mNET-seq (Nojima et al., 2015) to obtain single nucleotide resolution profiles was adapted to define the 5'end of the first read in each pair as well as its directionality. Exceptionally, for the POINT-seq DRB experiment, a spike-in SIRV-Set2 RNA was added before library preparation to allow comparison between control and DRB-treated cells. Here, reads were aligned against both SIRV-Set2 sequences available in the Lexogen website (version 170612a) and the reference human genome (GRCh38), using STAR software. Then, reads were counted using SAMtools (Li et al., 2009), considering their directionality based on SAM bitwise flag, and normalized as follows:

$$Normalized\ signal = \frac{HgR\ \times\ Sk}{10^6}$$

$HgR$ represents the number of reads aligned against the human genome from a particular region of interest, and Sk represents the total number of read counts aligned against the SIRV-Set 2 sequences. Division by $10^6$ was applied to improve readability. To evaluate experimental reproducibility, 2-3 biological replicates were generated. Read counts per replicate for each expressed protein-coding gene were obtained using *BedTools coverage* (Quinlan and Hall, 2010), requiring the same strand for the read and gene (-*s*). Splicing patterns were considered for POINT-seq (*-split*). Furthermore, *-counts* and *-sorted* parameters were added to the command. Spearman's rank-order test was then applied to discover the correlation between samples ($\rho$). ChrRNA-seq and pA+ RNA-seq were generated as part of GSE60358 and GSE86661; GSE33480, respectively. Their pre-processing was as for POINT-seq. Strand-specific CAGE from a previously published study (Chen et al., 2016) employed the same pre-processing as for POINT-5, where the 5'end of the original RNA was extracted from the entire sequenced read.

**POINT-nano pre-processing**

Nanopore raw signal fast5 files were basecalled using Guppy Basecalling 3.0.5 (Oxford Nanopore Technology Ltd.). NanoQC (De Coster et al., 2018) was used for a first evaluation of run sequencing quality. Since several samples were sequenced together, barcodes (NBD104/NBD114) were incorporated into the direct cDNA nanopore reads and identified with qcat. Extracting POINT-nano read directionality is critical, since it reveals transcript orientation. Thus, primer GAAGATAGAGCGACAGGCAAGT was searched for in reads using *regEx* Python package, applying the following rules: i<=3, d<=3, s<=3 and 1i+1d+1s <= 4. Only these reads were preserved with all others discarded since the Pol II position could not be determined. Barcode and primer sequences in validated reads were trimmed with Porechop, with --*discard_middle* mode on. Subsequently those were aligned with *minimap2* (Li, 2018) with *-ax splice* parameters. *Unmapped reads*, *not primary alignment* or *supplementary reads* were discarded using SAMtools bitwise flag 2308. To prevent contamination from non-authentic 3'ends due to oligodT priming on internal A rich sequences, reads with any T either in first 2 mapped bases or 3 out of 5 in the same region were ignored for downstream analyses. Finally, reads with 5'end soft clips longer than 50 bases were discarded. Pol II location was determined by the left most coordinate of the read, extracted with pysam package. Classification of Pol II position over the classes *TSS*, *Exon*, *Intron*, *SS*, *TES* and *Post-TES* was performed using *BedTools Intersect* after extracting these regions in a BED file format for expressed genes. While *TSS* and *TES* classes were classified over a 50 base region in both directions, SS was defined over a 10 base region. Post-TES region was determined by [TES+50, TES+550].

**Identification of expressed genes**

To determine the expressed genes in HeLa S3 and HCT116 cells, strand-specific pA+ RNA-seq data from previously published studies (Hela S3: GSE86661; HCT116: GSE33480) was used. Adaptors were trimmed with TrimGalore using the same parameters as in POINT-seq pre-processing. Then, *Kallisto* (Bray et al., 2016) mapped the reads against the human transcriptome (Ensembl v90), and TPM measurement for each transcription unit (TU) from the output was acquired. The transcript with highest TPM was selected per gene. Genes having no transcript with TPM higher than 4 were discarded. Moreover, filtered TUs must have protein-coding tag as a biotype, which was extracted from Ensembl GTF file version 90. To better detect signal levels from POINT technology in different experiments, overlapping TUs were excluded. To do this, an extra window of 500 bases upstream and 2000 bases downstream of each TU was added. A final number of 6341 and 5028 genes was identified as expressed in HeLa S3 and HCT116, respectively. Exceptionally metagene side windows of 2 kb upstream and 7 kb downstream were employed to exclude overlapping TUs. This led to 4546 and 5028 genes for HeLa S3 and HCT116, respectively. U snRNAs https://www.genenames.org/cgi-bin/genegroup/download?id=849&type=node, and histone genes https://www.genenames.org/cgi-bin/genegroup/download?id=864&type=branch, are described as part of *HUGO Gene Nomenclature Committee* platform. TUs from these classes without POINT reads were excluded from analyses.

#### Metagene Analysis

Metagenes were used to represent average Pol II, RNA 5'end and RNA distribution levels along genes and their flanks. To generate them, pre-processed BAM files split by forward and reverse strand were used as input for deepTools *bamCoverage* function. A black list of regions for hg38 assembly has been described (https://www.encodeproject.org/files/ENCFF419RSJ/), and all the reads from these were discarded. Also, RPKM normalisation was applied. Then, all expressed genes were scaled to have their TSS and TES overlapping, in a bin size of 10 bases, using the function deepTools *computeMatrix* (Ramirez et al., 2014) in *scale-regions* mode. Signal was captured not only from the gene, but also from 2 and 7 kb upstream and downstream flanking regions, respectively. In a final step, signal was displayed using the deepTools *computeMatrixOperation*, which generated one final table per strand containing the signal for each gene per bin. These tables were the input for a second phase of processing in R script. Here, genes with no signal in all bins and all conditions were removed as well as 1% of genes with the highest and lowest signals, as these could contribute to false average profiles. Finally, ggplot was used to graphically create the metagene.

#### Heatmaps

Heatmaps were employed to allow closer scrutiny of POINT-5 or CAGE-seq 5'end signal for each region individually. Thus, three different heatmap groups were built: mRNA-mRNA, PROMPT-mRNA and eRNA-eRNA, which were extracted from a previously published study (Chen et al., 2016). As with metagenes, read counts were captured from those datasets in HeLa S3 cells, using deepTools *bamCoverage* function, followed by deepTools *computeMatrix,* but in *reference-point* mode, preserving a bin size of 10 bases. Midpoint was obtained as the equidistant coordinate to the TSSs of both TUs. Regions with no signal in all conditions were removed. Obtained signal from minus strand was multiplied by -1, and then summed to the signal from the positive strand. Then, it was scaled using *rescale* function from *scales* R package. Lastly, *geom_raster* from ggplot package was employed to create the heatmap plot.

#### Cleavage ratio

Cleavage Ratio and Termination index were computed for POINT-5 data in a single-nucleotide basis. Cleavage Ratio was defined as:

$$CR = \frac{TES}{TSS + TES}$$

where TSS is the read counts per kilobase per million reads (FPKM) in the interval [TSS-50, TSS+50] and TES the FPKM for [TES-50, TES+50].

**Termination index**

The Termination Index was given by:

$$TI = log2\left(\frac{\frac{[TES,\ TES + 2000]_{counts}}{2000}}{\frac{GB_{counts}}{length_{GB}}}\right)$$

where GB stands for genome body and $length_{GB}$ is the number of bp between TSS and TES coordinates.

**Premature Transcription Termination Analysis**

Premature transcription terminated (PTT) genes were identified by comparison between control DMSO and PlaB POINT-seq data, using a simulation basis approach. Each gene was divided into 10bins. FPKM of each bin for DMSO was measured ($FPKM_{real}$). All reads overlapping the gene under PlaB condition were also randomly sampled, and the FPKM for each bin was measured ($FPKM_{simulation}$). This simulation was repeated 5000 times for each gene. To obtain more robust results and eliminate potential false negative hits, $FPKM_{simulation}$ was divided by a 3.5, value discovered by manual curation, giving rise to $FPKM_{T\_simulation}$. For each simulation, $FPKM_{T\_simulation}$ was compared with $FPKM_{real}$ for each bin and counted for how many times $FPKM_{T\_simulation}$ was lower than $FPKM_{real}$. Starting from the first bin for gene, whenever $FPKM_{T\_simulation}$ was found lower than $FPKM_{real}$ in at least 90% of the simulations for one bin, the search ended. This led to the conclusion that PTT occurs in that bin region. PTT genes were classified into Early (E), Middle (M), Late (L), according to where the bins were identified. Thus, Early (E) PTT occurred in the first 3bins, and Middle (M) and Late (L) PTT, occurred between 4-6bins and 7-9bins, respectively. If PTT was ascribed to the last bin or was not found for any of the bins then it was considered non-PTT and labelled NC.

**Splicing analyses with POINT technology**

POINT-nano and POINT-seq data were used to dissect splicing kinetics. Only internal introns were considered for these analyses. Thus, genes with less than 3 introns were discarded, as well as first and last introns from genes with higher intronic complexity. Intron and exon sizes were extracted from Ensembl annotations GTF files for expressed genes previously identified. The 25% shortest and longest features only were taken to perform splicing comparative analyses. Two-sided Mann-Whitney test was deduced to obtain their significance, followed by a p values adjustment using Holm method. Furthermore, SS strength score was measured with *MaxEntScan* (Yeo and Burge, 2004) using default parameters. Several filters and transformations were applied to POINT-nano reads to reveal their splicing status per overlapped intron. Firstly, reads must span 10 bases from 3'SS to the downstream exon. This overlap was validated with *BedTools intersect*, with prior transformation of reads from BAM format to BED format using *BedTools bamtobed* in split mode (-split). A full overlap (-F 1) and a shared strand between read and exon (-s) was required here. Then, two different windows of 10 bases were created in addition to downstream 3'SS, upstream 3'SS and 5'SS, to analyse splicing patterns. Reads were considered unspliced when detected in the upstream 3'SS window. All reads which were not detected in this window, but only in upstream 5'SS, were considered spliced. Thus, the splicing fraction denotes the number of reads found as spliced divided by spliced and unspliced reads. Importantly, reads spanned over several genes were discarded. Only introns with their 3'SS 1500 bp upstream to Pol II were considered, except for **Figures 6** and **S6** were a distance of 3500 bp was accepted. Additionally, only the latest fully transcribed intron was considered, according to Pol II position. Exceptionally for **Figure 6F**, this was repeated for the 2 or 3 latest transcribed introns. Splicing status of POINT-seq reads for each intron was obtained as for POINT-nano, but by use of 5bp windows. For introns with more than 10 spliced and/or unspliced reads, the splicing fraction was measured with all the POINT-seq splicing analyses performed. Distance to TES from 3'SS was extracted from Ensembl human reference annotation, by subtracting its coordinates and assuming that the TES location is the end coordinate of annotated genes.

**Alternative splicing events and cassette cases identification**

Constitutive and alternative splicing event classifications were obtained from Ensembl annotations, as previously described (Nojima et al., 2018). For cases of cassette exon identification, previously published pA+ RNA-seq data from HeLa S3 cells (GSE86661) was analysed using *vast-tools (https://doi.org/10.1101/gr.220962.117)*. Exon skipping events were isolated, and exons were considered included or excluded when the Ψ value was higher than 0.75 or lower than 0.25, respectively.

**Signal extraction and data visualization**

Read Coverage and gene annotation manipulations were performed with *BedTools*. BAM files were split by strand with SAMtools according their bitwise flags. In POINT-seq data, forward oriented strand had 83 and 163 flags associated while reverse oriented strand had 99 and 147 flags associated. Oppositely, 99 and 147 flags for POINT-5 correspond to the forward strand, while 83 and 163 to the reverse strand. For POINT-nano, 0 and 16 flags were used to call forward and reverse strand reads, respectively. Data was visualized applying *genomeCoverageBed* function of *BedTools* to each strand independently. Trackhubs in the UCSC browser were created by employing the UCSC bedGraphToBigWig tool (Kent et al., 2002).

**Reads quantification**

Reads were counted for regions of interest with *BedTools Intersect*, using post normalization to library size and gene length using *read counts per kilobase per million reads* (FPKM).

**P-values and significance tests**

Significance between control and treatment condition was obtained using a two-sided Mann-Whitney test, followed by a p values adjustment using the Holm method. For multiple samples one-way ANOVA comparison was tested, followed by a post-hoc analysis using Turkey’s test.

### SUPPLEMENTARY FIGURE LEGENDS

**Figure S1. Additional notes of POINT-seq and POINT-5 methods (related to Figure 1)**

(A) Western blot of supernatant from DNase-digested HeLa chromatin treated with indicated concentration of Empigen in NUN2. Antibodies used against Pol II CTD (CMA601) and histone H3.

(B) DNA fragments in supernatant of DNase-digested HeLa chromatin treated with 3% Empigen in NUN2 for indicated incubation times.

(C) Western blot of supernatant from DNase-digested HeLa chromatin (input) and IP with Pol II CTD antibody. Antibodies for western blot were used against Pol II CTD (CMA601) and histone H3.

(D) Tapestation image of POINT RNA size distribution. Digested chromatin fraction as input (blue), IP with mouse IgG (red), and IP with Pol II CTD antibody CMA601 (green).

(E) (Left) *TARS* of example of POINT-seq signals and (Right) quantification of POINT-seq metagene (n=3067) signals from DMSO or DRB (4 hr)-treated HeLa cells.

(F) Heat map for reproducibility between three biological replicates of POINT-seq in HeLa cells.

(G) Scatter plot for reproducibility between two biological replicates of POINT-5 in HeLa cells.

(H) Example view of POINT-5 and POINT-seq signals for mRNA-mRNA pairs in HeLa cells.

(I) Example view of POINT-5 and POINT-seq signals for eRNA-eRNA pairs in HeLa cells.

(J) Heat maps of POINT-5 and CAGE (-/+ RRP40KD) for eRNA-eRNA pairs in HeLa cells.

**Figure S2. Duplicates of POINT-5 and POINT-seq gene profiles (related to Figure 2)**

(A-D) Example views of POINT-5 and POINT-seq signals on (A) Histone gene cluster, (B) *CTDSP1* as miR26B host gene, (C) Enhancer and unannotated lncRNA with pre-miRNAs, and (D) *RNU5A-1 and RNU4-2* as U snRNA genes. Sense strand, blue. Antisense strand, red. S and C are represented as transcription start and RNA cleavage sites, respectively.

**Figure S3. Metagene profile and example genes of POINT-seq and POINT-5 in Xrn2 and CPSF73-depleted HCT116 cells (related to Figure 3)**

(A) Western blot of whole cell extract from HeLa and HCT116 Xrn2-AID (-/+ IAA, 4 hr). Antibodies against Xrn2 and Tubulin. Non-specific (n-sp) bands from Xrn2 antibody indicated by arrow heads.

(B and C) Metagene profile of POINT-seq (B) or POINT-5 (C) on normalised protein coding (PC) genes in HCT116 Xrn2-AID (-/+ IAA, 4 hr).

(D) Ratio of TES and TSS on PC genes in HeLa and HCT116 Xrn2-AID (- IAA) cells.

(E) *HIST1H4E* and *RNU1* as examples of POINT-5 and POINT-seq on histone and UsnRNA genes in HCT116 Xrn2-AID (-/+ IAA, 4 hr).

(F) *MIR17HG* and *VEZT* as examples of POINT-5 and POINT-seq on miRNA host genes in HCT116 Xrn2-AID (-/+ IAA, 4 hr).

(G) *JARID2* as example of POINT-5 and POINT-seq on PC gene in HCT116 Xrn2-AID (-/+ IAA, 4 hr) and *in vitro* ExT treatment. POINT-5 peaks located downstream of PAS in HCT116 Xrn2-AID (+ IAA, 4 hr) cell are indicated as green asterisks. ExT-sensitive POINT-5 peaks located downstream of TSS2 indicated by red arrows.

(H) Western blot of whole cell extract of HeLa and HCT116 CPSF73-AID (-/+ IAA, 3 hrs). Antibodies against mAID and Tubulin.

(I and J) Metagene profile of (I) POINT-seq or (J) POINT-5 on normalised PC genes in HCT116 CPSF73-AID (-/+ IAA, 3 hr).

**Figure S4. Additional analysis of PlaB induced PTT (related to Figure 4)**

(A) Venn graph of PTT genes induced by U1 AMO (8 hr) or PlaB (4 hr) treatment in HeLa cells.

(B) *MTHFD1L* and *DHX9* as examples of 4-shU RNA-seq and POINT-seq signals.

(C) (Top) Schematic of CPSF73KD and PlaB treatment in HCT116 CPSF73-AID cells. (Bottom) Metagene profile of POINT-seq with PlaB (light blue) or CPSF73KD and PlaB (orange) in normalized region from TSS-2kbp to TES+7kbp. Arrow indicates a starting point of PTT defect induced by CPSF73KD.

(D) *GPATCH2* as example of POINT-seq for indicated treatments.

**Figure S5. Additional characterization of POINT-nano (related to Figure 5)**

(A) Tapestation image of size distribution of *in vitro* polyadenylated POINT RNA.

(B and C) Consensus sequences of POINT-nano 5'end reads (B) before or (C) after removal of internal poly A tracts (reverse complement).

(D) Scatter plot for reproducibility between two biological replicates of POINT-nano signals in HeLa cells.

(E) Gene coverage of POINT-nano reads

(F) Pie charts of mNET-seq and POINT-nano RNA 3'ends for positions of protein coding genes.

(G) *ID1* as example of POINT-5, POINT-seq, and POINT-nano signals. Spliced (red), unspliced (blue), other (grey) reads.

**Figure S6. Additional bioinformatics on co-transcriptional splicing (related to Figure 6)**

(A) Splicing fraction of POINT-nano reads with Pol II located at 3'SS+10 to +49 bp in HeLa cells. Spliced (red), Unspliced (blue).

(B) Splicing fraction of POINT-nano reads with Pol II located downstream of 5'SS. Untreated, DMSO and PlaB-treated HeLa cell are shown as black, orange and blue dashed lines.

(C) Effect of 3'SS and 5'SS scores on splicing fraction in POINT-seq analyses.

(D) Effect of 3'SS and 5'SS scores (high 10% or low 10%) on splicing fraction in POINT-nano analyses with Pol II in exon and intron.

(E) Splicing fraction of POINT-seq signals for alternative splicing (AS) or constitutive splicing (CS).

(F) (Top) Cassette exon splicing events classified based on exon inclusion percentage using pA+ RNA-seq data. (Bottom) Number of exon inclusion (red) and exclusion (grey) events in POINT-nano analyses profiled for high or low 25% exon inclusion event categories.

(G) Exon inclusion or exclusion levels in POINT-nano analyses with Pol II located downstream of 3'SS in intron 2. High 25% and low 25% of exon inclusion events are shown as red and black lines, respectively. Numbers of splicing events are indicated.

(H) Splicing fraction for last intron removal in POINT-nano analyses. Pol II positions indicated below.

**Figure S7. Additional bioinformatics on TU length effect on co-transcriptional splicing (related to Figure 7)**

(A-D) Effect of (A and B) exon or (C and D) intron sizes (long 25%, red or short 25%, black) on splicing fraction in POINT-nano analyses. Pol II is located downstream of (A and C) 3'SS (5'SS+0~1.5Kbp) or (B and D) 5'SS (5'SS+0~1Kbp).

(E) Quantification of effect of exon (top) or intron (bottom) sizes on splicing fraction POINT-nano signals. Pol II positions are indicated at bottom.

(F) Splicing fraction of POINT-seq signals in HeLa cells. Intron positions and distance from 3'ss to TES (kbp) are indicated at top and bottom, respectively.

(G) Splicing fraction of POINT-seq signals in HeLa cells. Distance from 3'ss to TES (kbp) is indicated below. AS (yellow), CS (blue).

(H) *HNRNPA0* as example of POINT-seq in HCT116 CPSF73-AID cells (-/+ IAA). CPSF73KD induced read-in *KLHL3* RNA. Exonic signals are pointed by red arrows.

(I) Number of introns with splicing ratio of POINT-seq signals in HCT116 CPSF73-AID cells (-/+ IAA). Control (blue), CPSF73KD (light red).

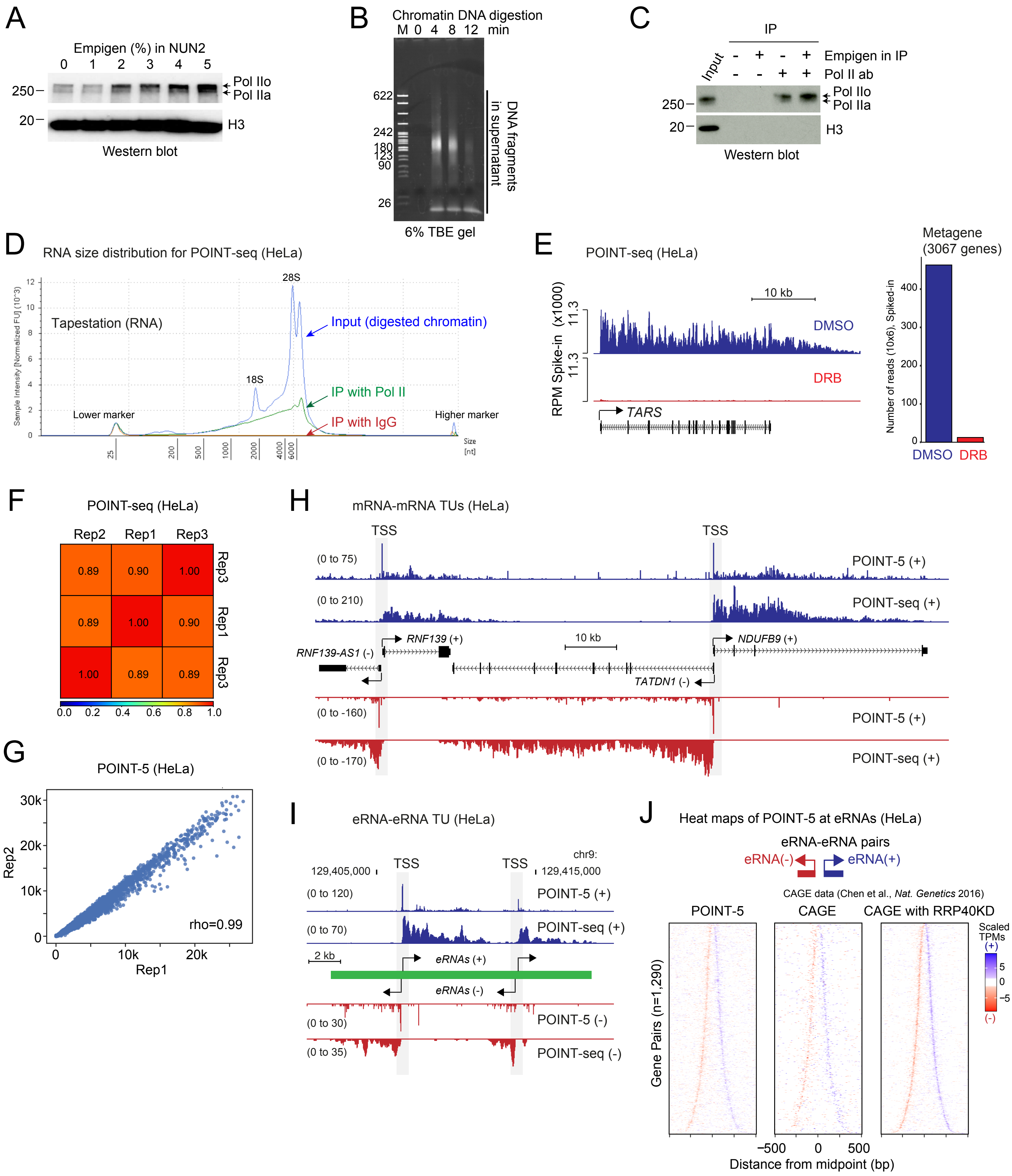
A
Empigen (%) in NUN2
0 1 2 3 4 5
250
20
Pol IIo
Pol IIa
H3
Western blot
B
Chromatin DNA digestion
M 0 4 8 12 min
622
242
180
123
90
26
DNA fragments in supernatant
6% TBE gel
C
IP
Input
Empigen in IP
Pol II ab
250
20
Pol IIo
Pol IIa
H3
Western blot
D
RNA size distribution for POINT-seq (HeLa)
Tapestation (RNA)
28S
18S
Input (digested chromatin)
IP with Pol II
IP with IgG
Lower marker
Higher marker
Sample Intensity [Normalized FU] (10^3)
Size [nt]
E
POINT-seq (HeLa)
RPM Spike-in (x1000)
10 kb
11.3
DMSO
DRB
TARS
Metagene (3067 genes)
Number of reads (10x6), Spiked-in
DMSO
DRB
F
POINT-seq (HeLa)
Rep2 Rep1 Rep3
G
POINT-5 (HeLa)
Rep2
Rep1
rho=0.99
H
mRNA-mRNA TUs (HeLa)
TSS
TSS
(0 to 75)
POINT-5 (+)
(0 to 210)
POINT-seq (+)
RNF139 (+)
10 kb
NDUFB9 (+)
RNF139-AS1 (-)
TATDN1 (-)
(0 to -160)
POINT-5 (+)
(0 to -170)
POINT-seq (+)
I
eRNA-eRNA TU (HeLa)
chr9:
129,405,000
TSS
TSS
129,415,000
(0 to 120)
POINT-5 (+)
(0 to 70)
POINT-seq (+)
2 kb
eRNAs (+)
eRNAs (-)
(0 to 30)
POINT-5 (-)
(0 to 35)
POINT-seq (-)
J
Heat maps of POINT-5 at eRNAs (HeLa)
eRNA-eRNA pairs
eRNA(-)
eRNA(+)
CAGE data (Chen et al., Nat. Genetics 2016)
POINT-5
CAGE
CAGE with RRP40KD
Gene Pairs (n=1,290)
Scaled TPMs
Distance from midpoint (bp)


Figure S1

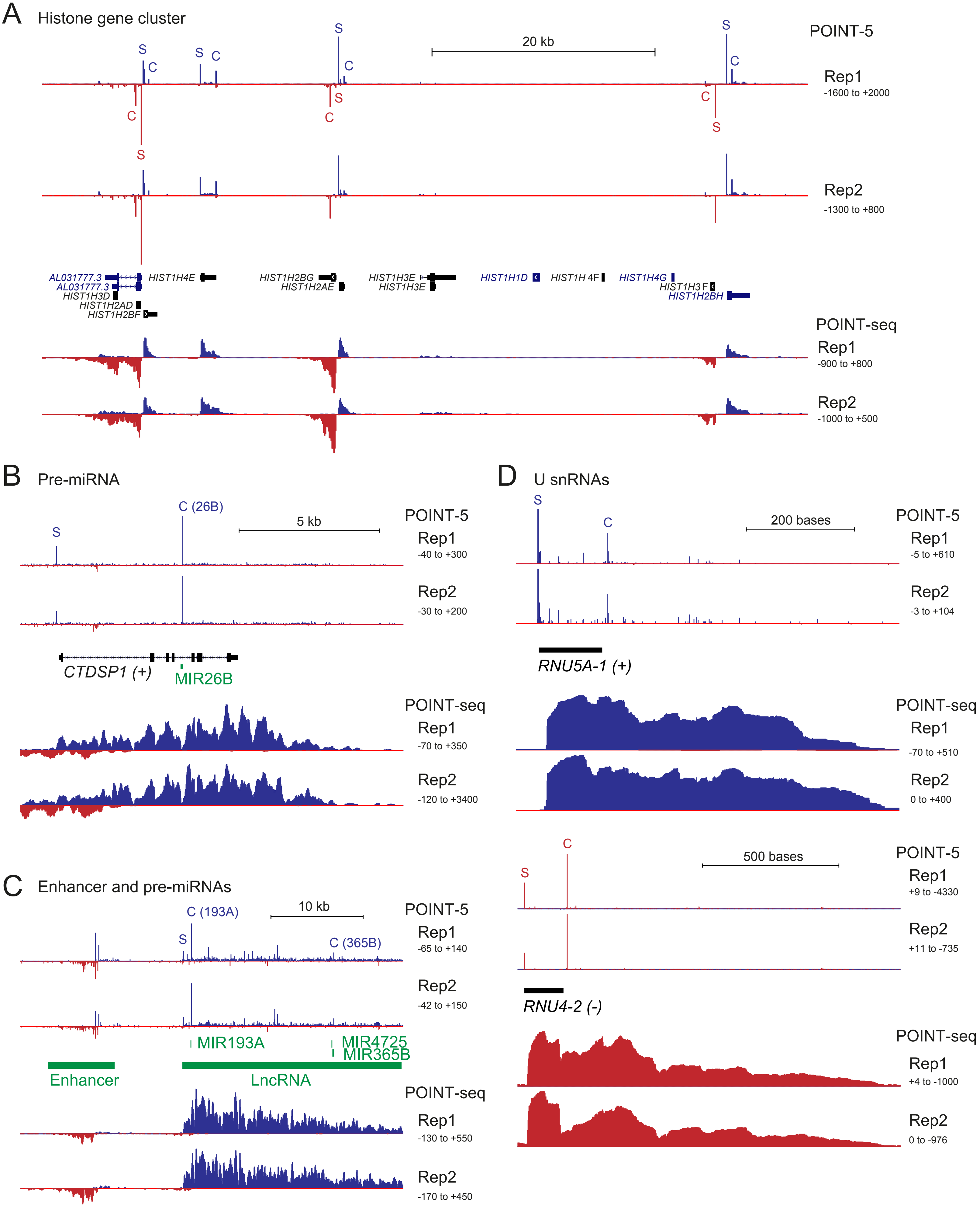
A
Histone gene cluster
POINT-5
20 kb
S
C
Rep1
-1600 to +2000
Rep2
-1300 to +800
AL031777.3
AL031777.3
HIST1H3D
HIST1H2AD
HIST1H2BF
HIST1H4E
HIST1H2BG
HIST1H2AE
HIST1H3E
HIST1H3E
HIST1H1D
HIST1H 4F
HIST1H4G
HIST1H3 F
HIST1H2BH
POINT-seq
Rep1
-900 to +800
Rep2
-1000 to +500
B
Pre-miRNA
C (26B)
5 kb
POINT-5
Rep1
-40 to +300
Rep2
-30 to +200
CTDSP1 (+)
MIR26B
POINT-seq
Rep1
-70 to +350
Rep2
-120 to +3400
C
Enhancer and pre-miRNAs
C (193A)
10 kb
C (365B)
POINT-5
Rep1
-65 to +140
Rep2
-42 to +150
MIR193A
MIR4725
MIR365B
Enhancer
LncRNA
POINT-seq
Rep1
-130 to +550
Rep2
-170 to +450
D
U snRNAs
200 bases
POINT-5
Rep1
-5 to +610
Rep2
-3 to +104
RNU5A-1 (+)
POINT-seq
Rep1
-70 to +510
Rep2
0 to +400
500 bases
POINT-5
Rep1
+9 to -4330
Rep2
+11 to -735
RNU4-2 (-)
POINT-seq
Rep1
+4 to -1000
Rep2
0 to -976

Figure S2

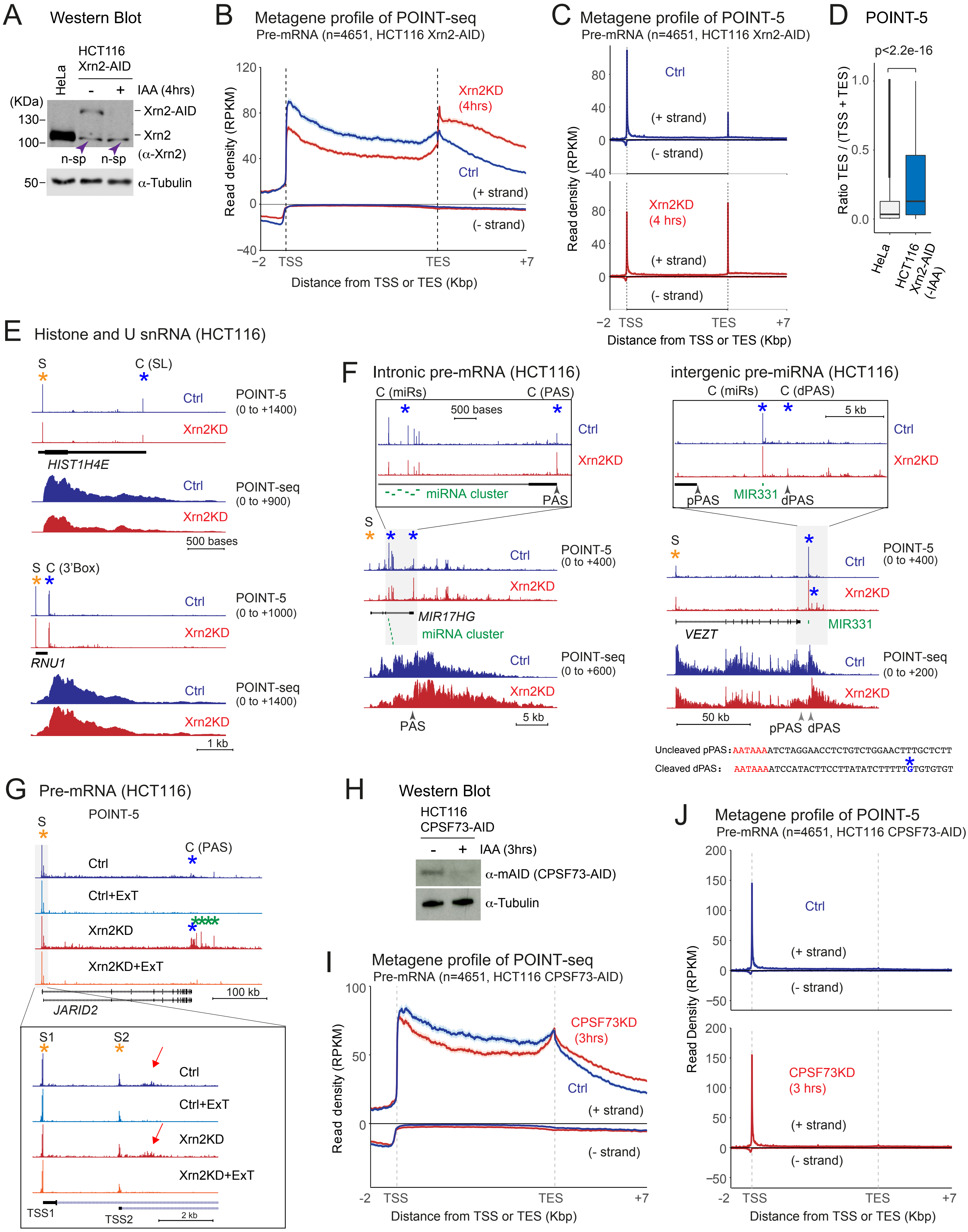

A Western Blot
HCT116 Xrn2-AID
HeLa
- + IAA (4hrs)
(KDa)
130–
100–
Xrn2-AID
Xrn2
(α-Xrn2)
n-sp n-sp
50–
α-Tubulin
B Metagene profile of POINT-seq
Pre-mRNA (n=4651, HCT116 Xrn2-AID)
Read density (RPKM)
Xrn2KD (4hrs)
Ctrl
(+ strand)
(- strand)
–2 TSS TES +7
Distance from TSS or TES (Kbp)
C Metagene profile of POINT-5
Pre-mRNA (n=4651, HCT116 Xrn2-AID)
Ctrl
(+ strand)
(- strand)
Xrn2KD (4 hrs)
Read density (RPKM)
–2 TSS TES +7
Distance from TSS or TES (Kbp)
D POINT-5
p<2.2e-16
Ratio TES / (TSS + TES)
HeLa
HCT116 Xrn2-AID (-IAA)
E Histone and U snRNA (HCT116)
S C (SL)
Ctrl
Xrn2KD
POINT-5 (0 to +1400)
HIST1H4E
POINT-seq (0 to +900)
500 bases
S C (3'Box)
POINT-5 (0 to +1000)
RNU1
POINT-seq (0 to +1400)
1 kb
F Intronic pre-mRNA (HCT116)
C (miRs) C (PAS)
500 bases
miRNA cluster
PAS
S
POINT-5 (0 to +400)
MIR17HG
miRNA cluster
POINT-seq (0 to +600)
PAS
5 kb
intergenic pre-miRNA (HCT116)
C (miRs) C (dPAS)
5 kb
pPAS MIR331 dPAS
POINT-5 (0 to +400)
MIR331
VEZT
POINT-seq (0 to +200)
50 kb
pPAS dPAS
Uncleaved pPAS: AATAAAATCTAGGAACCTCTGTCTGGAACTTTGCTCTT
Cleaved dPAS: AATAAAATCCATACTTCCTTATATCTTTTTGTGTGTGT
G Pre-mRNA (HCT116)
POINT-5
S
C (PAS)
Ctrl
Ctrl+ExT
Xrn2KD
Xrn2KD+ExT
100 kb
JARID2
S1 S2
TSS1 TSS2
2 kb
H Western Blot
HCT116 CPSF73-AID
- + IAA (3hrs)
α-mAID (CPSF73-AID)
α-Tubulin
I Metagene profile of POINT-seq
Pre-mRNA (n=4651, HCT116 CPSF73-AID)
Read density (RPKM)
CPSF73KD (3hrs)
Ctrl
(+ strand)
(- strand)
-2 TSS TES +7
Distance from TSS or TES (Kbp)
J Metagene profile of POINT-5
Pre-mRNA (n=4651, HCT116 CPSF73-AID)
Read Density (RPKM)
Ctrl
CPSF73KD (3 hrs)
(+ strand)
(- strand)
-2 TSS TES +7
Distance from TSS or TES (Kbp)


Figure S3

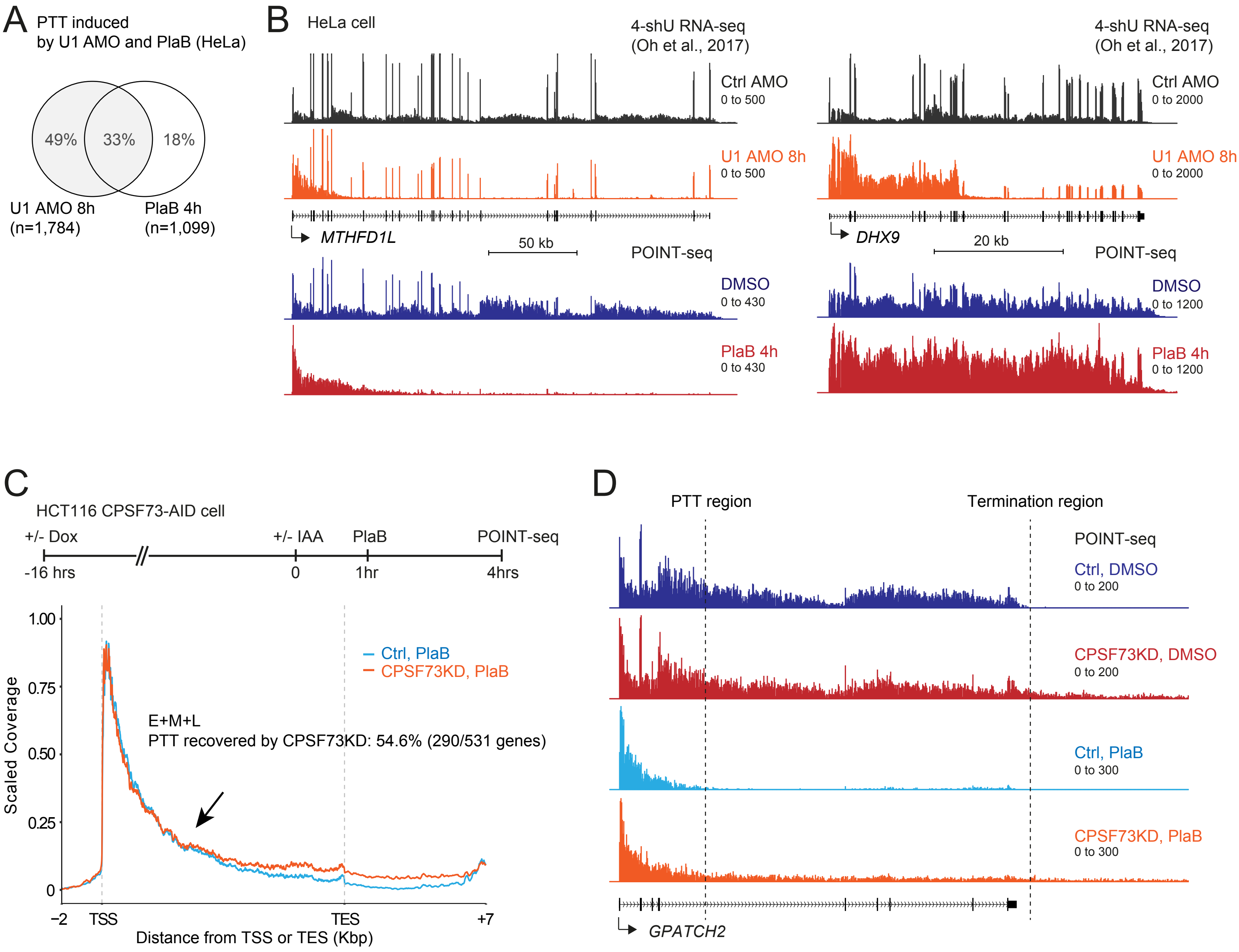
A
PTT induced by U1 AMO and PlaB (HeLa)
49%
33%
18%
U1 AMO 8h (n=1,784)
PlaB 4h (n=1,099)
B
HeLa cell
4-shU RNA-seq (Oh et al., 2017)
Ctrl AMO
0 to 500
U1 AMO 8h
0 to 500
MTHFD1L
50 kb
POINT-seq
DMSO
0 to 430
PlaB 4h
0 to 430
4-shU RNA-seq (Oh et al., 2017)
Ctrl AMO
0 to 2000
U1 AMO 8h
0 to 2000
DHX9
20 kb
POINT-seq
DMSO
0 to 1200
PlaB 4h
0 to 1200
C
HCT116 CPSF73-AID cell
+/- Dox
+/- IAA
PlaB
POINT-seq
-16 hrs
0
1hr
4hrs
Ctrl, PlaB
CPSF73KD, PlaB
E+M+L
PTT recovered by CPSF73KD: 54.6% (290/531 genes)
Scaled Coverage
1.00
0.75
0.50
0.25
0
−2
TSS
TES
+7
Distance from TSS or TES (Kbp)
D
PTT region
Termination region
POINT-seq
Ctrl, DMSO
0 to 200
CPSF73KD, DMSO
0 to 200
Ctrl, PlaB
0 to 300
CPSF73KD, PlaB
0 to 300
GPATCH2

Figure S4

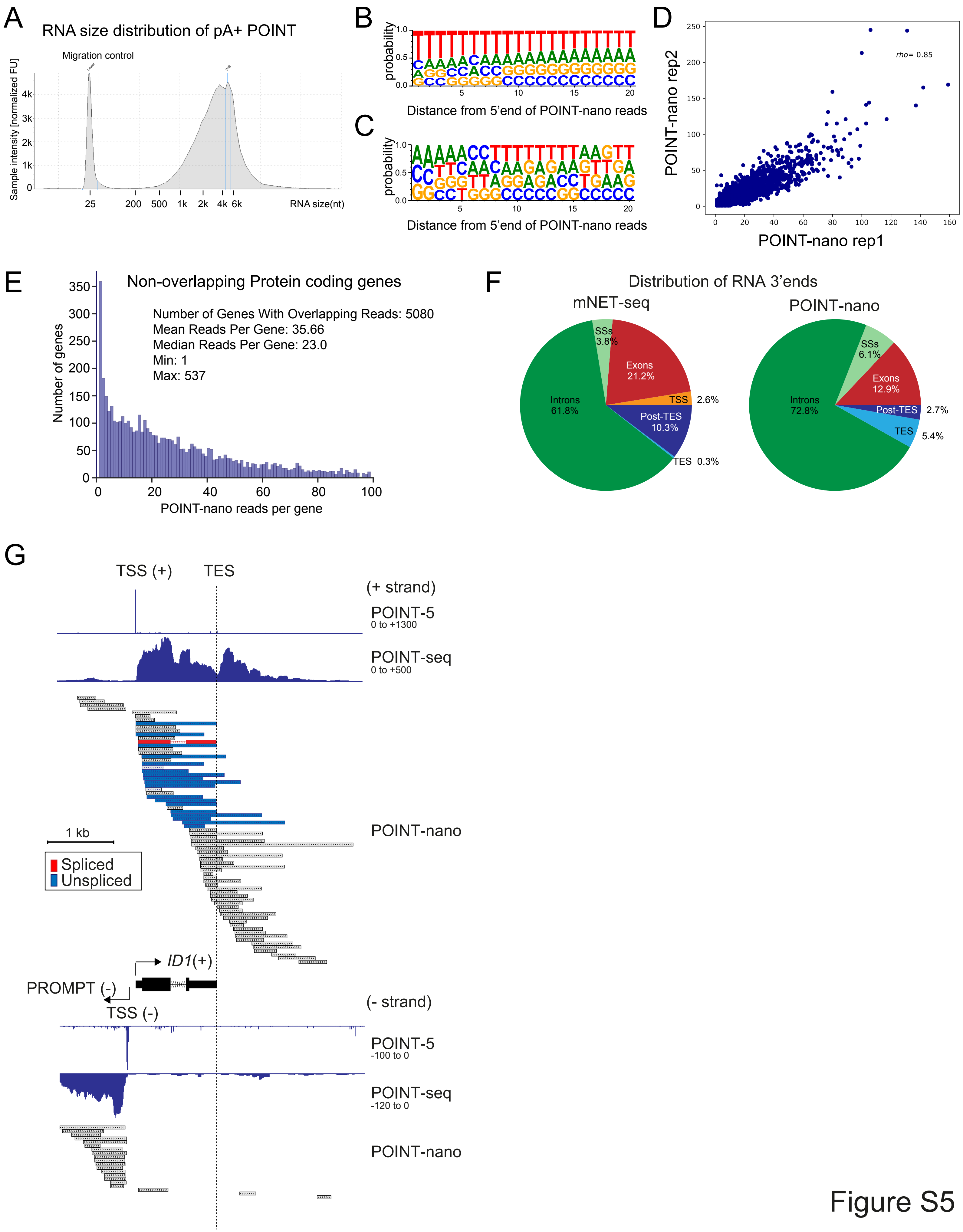
A
RNA size distribution of pA+ POINT
Migration control
Sample intensity [normalized FU]
RNA size(nt)
B
probability
Distance from 5'end of POINT-nano reads
C
probability
Distance from 5'end of POINT-nano reads
D
POINT-nano rep2
POINT-nano rep1
rho= 0.85
E
Non-overlapping Protein coding genes
Number of Genes With Overlapping Reads: 5080
Mean Reads Per Gene: 35.66
Median Reads Per Gene: 23.0
Min: 1
Max: 537
Number of genes
POINT-nano reads per gene
F
Distribution of RNA 3'ends
mNET-seq
SSs 3.8%
Exons 21.2%
TSS 2.6%
Post-TES 10.3%
TES 0.3%
Introns 61.8%
POINT-nano
SSs 6.1%
Exons 12.9%
Post-TES 2.7%
TES 5.4%
Introns 72.8%
G
TSS (+)
TES
(+ strand)
POINT-5
0 to +1300
POINT-seq
0 to +500
POINT-nano
1 kb
Spliced
Unspliced
ID1(+)
PROMPT (-)
TSS (-)
(- strand)
POINT-5
-100 to 0
POINT-seq
-120 to 0
POINT-nano

Figure S5

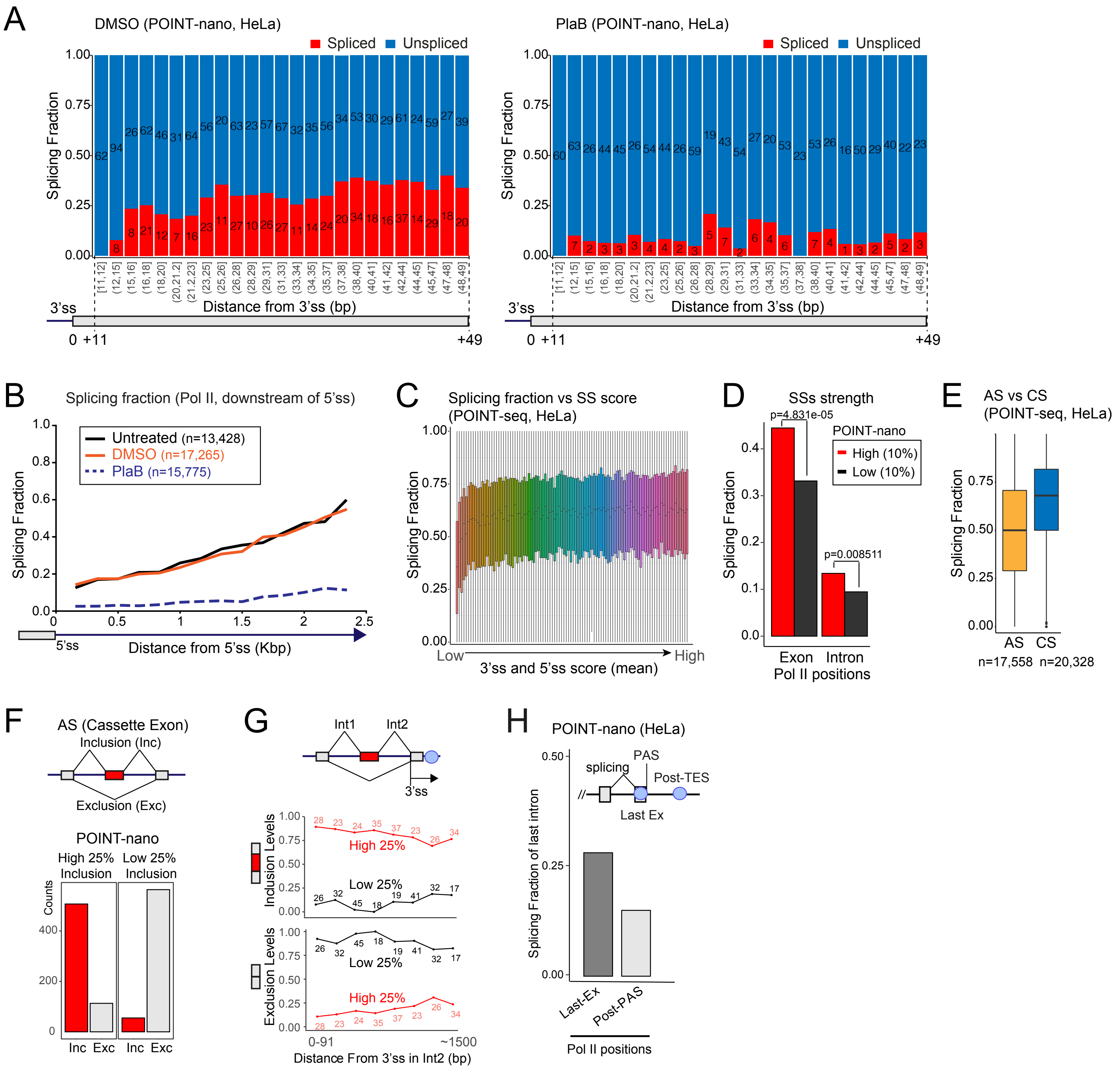
A
DMSO (POINT-nano, HeLa)
Spliced
Unspliced
Splicing Fraction
Distance from 3'ss (bp)
3'ss
0 +11
+49
PlaB (POINT-nano, HeLa)
B
Splicing fraction (Pol II, downstream of 5'ss)
Untreated (n=13,428)
DMSO (n=17,265)
PlaB (n=15,775)
5'ss
Distance from 5'ss (Kbp)
C
Splicing fraction vs SS score (POINT-seq, HeLa)
Low
High
3'ss and 5'ss score (mean)
D
SSs strength
p=4.831e-05
p=0.008511
POINT-nano
High (10%)
Low (10%)
Exon
Intron
Pol II positions
E
AS vs CS (POINT-seq, HeLa)
AS
CS
n=17,558
n=20,328
F
AS (Cassette Exon)
Inclusion (Inc)
Exclusion (Exc)
POINT-nano
High 25% Inclusion
Low 25% Inclusion
Counts
Inc Exc
Inc Exc
G
Int1
Int2
3'ss
Inclusion Levels
High 25%
Low 25%
Exclusion Levels
0-91
~1500
Distance From 3'ss in Int2 (bp)
H
POINT-nano (HeLa)
splicing
PAS
Post-TES
Last Ex
Splicing Fraction of last intron
Last-Ex
Post-PAS
Pol II positions

Figure S6

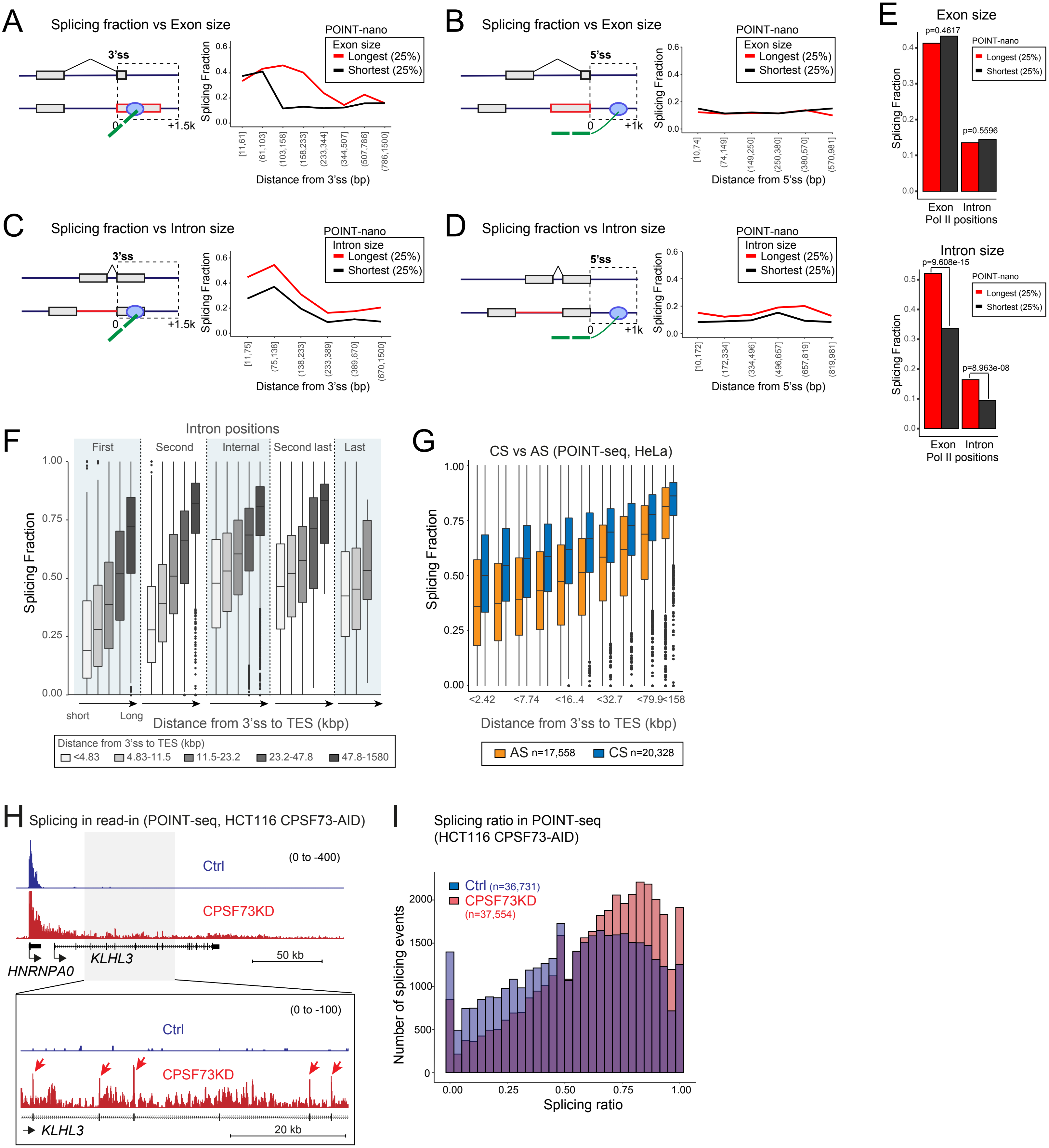
A
Splicing fraction vs Exon size
3'ss
0
+1.5k
POINT-nano
Exon size
Longest (25%)
Shortest (25%)
Splicing Fraction
Distance from 3'ss (bp)
B
Splicing fraction vs Exon size
5'ss
0
+1k
POINT-nano
Exon size
Longest (25%)
Shortest (25%)
Splicing Fraction
Distance from 5'ss (bp)
C
Splicing fraction vs Intron size
3'ss
0
+1.5k
POINT-nano
Intron size
Longest (25%)
Shortest (25%)
Splicing Fraction
Distance from 3'ss (bp)
D
Splicing fraction vs Intron size
5'ss
0
+1k
POINT-nano
Intron size
Longest (25%)
Shortest (25%)
Splicing Fraction
Distance from 5'ss (bp)
E
Exon size
p=0.4617
p=0.5596
POINT-nano
Longest (25%)
Shortest (25%)
Splicing Fraction
Exon
Intron
Pol II positions
Intron size
p=9.608e-15
p=8.963e-08
POINT-nano
Longest (25%)
Shortest (25%)
Splicing Fraction
Exon
Intron
Pol II positions
F
Intron positions
First
Second
Internal
Second last
Last
Splicing Fraction
short
Long
Distance from 3'ss to TES (kbp)
Distance from 3'ss to TES (kbp)
<4.83
4.83-11.5
11.5-23.2
23.2-47.8
47.8-1580
G
CS vs AS (POINT-seq, HeLa)
Splicing Fraction
Distance from 3'ss to TES (kbp)
AS n=17,558
CS n=20,328
H
Splicing in read-in (POINT-seq, HCT116 CPSF73-AID)
Ctrl
(0 to -400)
CPSF73KD
HNRNPA0
KLHL3
50 kb
(0 to -100)
Ctrl
CPSF73KD
KLHL3
20 kb
I
Splicing ratio in POINT-seq
(HCT116 CPSF73-AID)
Ctrl (n=36,731)
CPSF73KD
(n=37,554)
Number of splicing events
Splicing ratio

Figure S7
